# A Thalamocortical Cell-Adhesion Code Primes Sensory Cortical Specification

**DOI:** 10.1101/2025.11.17.688831

**Authors:** T. Guillamón-Vivancos, L. Puche-Aroca, L. Wang, I. Fuentes-Jurado, D. Vandael, D. Torres, M. Aníbal-Martínez, FJ. Martini, G. López-Bendito

## Abstract

The emergence of functional sensory modalities requires precise cortical arealization and appropriate thalamocortical targeting. Although early morphogen gradients set broad cortical territories, the mechanisms that specify sensory identity and guide modality-specific thalamocortical targeting remain unknown. Here, we identify an embryonic set of activity-independent cortical “protogenes”, prominently enriched for cell-adhesion molecules, that are differentially expressed between primary somatosensory (S1) and visual (V1) cortices prior to thalamic innervation. These adhesion programs are selectively localized to layer 4, the main thalamo-recipient layer, and strikingly mirror the adhesion profiles of their corresponding thalamic nuclei, suggesting partner recognition. Disrupting thalamic activity alters modality-specific spontaneous cortical dynamics and the postnatal expression of another set of modality-specific genes, indicating a modulatory role for thalamic activity in cortical identity. These findings support a two-step model in which cortical identity is primed by adhesion codes driving modality-specific thalamocortical targeting, and later refined by patterned thalamic activity to establish functional cortical modalities.

---

The mammalian neocortex is parcellated into specialized sensory areas, each dedicated to processing a modality specific thalamic input. Classical models posit that early morphogen gradients initiate cortical patterning, establishing the broad areas^1–5^ that are subsequently influenced by highly specific thalamic inputs. Gradients of transcription factors (TF), such as *Emx2* and *Pax6,* generate coarse rostrocaudal and mediolateral identities and influence thalamocortical axon guidance^6–9^. In parallel, human developmental studies at single-cell resolution have revealed topographic transcriptional signatures across the embryonic cortical sheet^10–12^.

Although these intrinsic programs establish broad cortical territories, definitive cortical identity also requires thalamic input. During early development, thalamocortical afferents not only provide patterned spontaneous activity to the cortex^13–15^ but also influence the transcriptional programs of postsynaptic cortical neurons^16,17^. Experimental perturbations of thalamic activity disrupt the size and organization of sensory maps^13–15^, and depletion or misrouting of thalamic projections can alter cortical areal boundaries^18,19^. These findings suggest that cortical modality identities, such as somatosensory versus visual cortex, arise from a combination of intrinsic and thalamus-dependent mechanisms. Yet this framework still overlooks a fundamental issue: how distinct thalamic nuclei reliably recognize and innervate their appropriate cortical targets.

Here, we examine how cortical areas acquire molecular and functional modality identities through the interplay of intrinsic genetic programs and thalamic activity, and how these identities may, in turn, guide modality-specific thalamocortical targeting. Using single nucleus RNA sequencing (snRNA-seq), spatial transcriptomics and *in vivo* calcium imaging, we uncover a set of embryonic, activity-independent “protogenes” encoding cell-adhesion molecules that are differentially expressed between primary somatosensory (S1) and visual (V1) cortices before thalamocortical synaptogenesis. These adhesion signatures are concentrated in layer 4 (L4), the principal thalamo-recipient layer, and remarkably mirror the adhesion profiles of their respective thalamic nuclei, ventral posteromedial (VPM) and dorsal lateral geniculate (dLG), potentially enabling homophilic thalamo-cortical interactions. Subsequently, patterned thalamic activity refines, but does not establish, these programs, indicating a two-step mechanism in which intrinsic adhesion codes prime thalamocortical targeting and activity consolidates functional modality identity. Together, our findings reveal a previously unrecognized molecular logic for the assembly of thalamocortical circuits and provide a new conceptual framework for understanding how sensory modality identity emerges within the developing neocortex.

## A modality-specific transcriptional program emerges in layer 4

By postnatal day 4 (P4), thalamocortical axons delineate the visual and somatosensory territories histologically^20^ and several genes are differentially expressed in these emerging sensory areas when compared to rostral-motor cortex^21–23^. However, a direct comprehensive comparison of S1 and V1 transcriptomes at this stage has been lacking. We therefore profiled S1 and V1 at P4 using snRNA-seq (Fig. 1a). All expected cortical cell types were identified, with modality differences mostly confined to glutamatergic neurons (Fig. S1), consistent with findings in human cortex^10,11^.

**Fig. 1.**
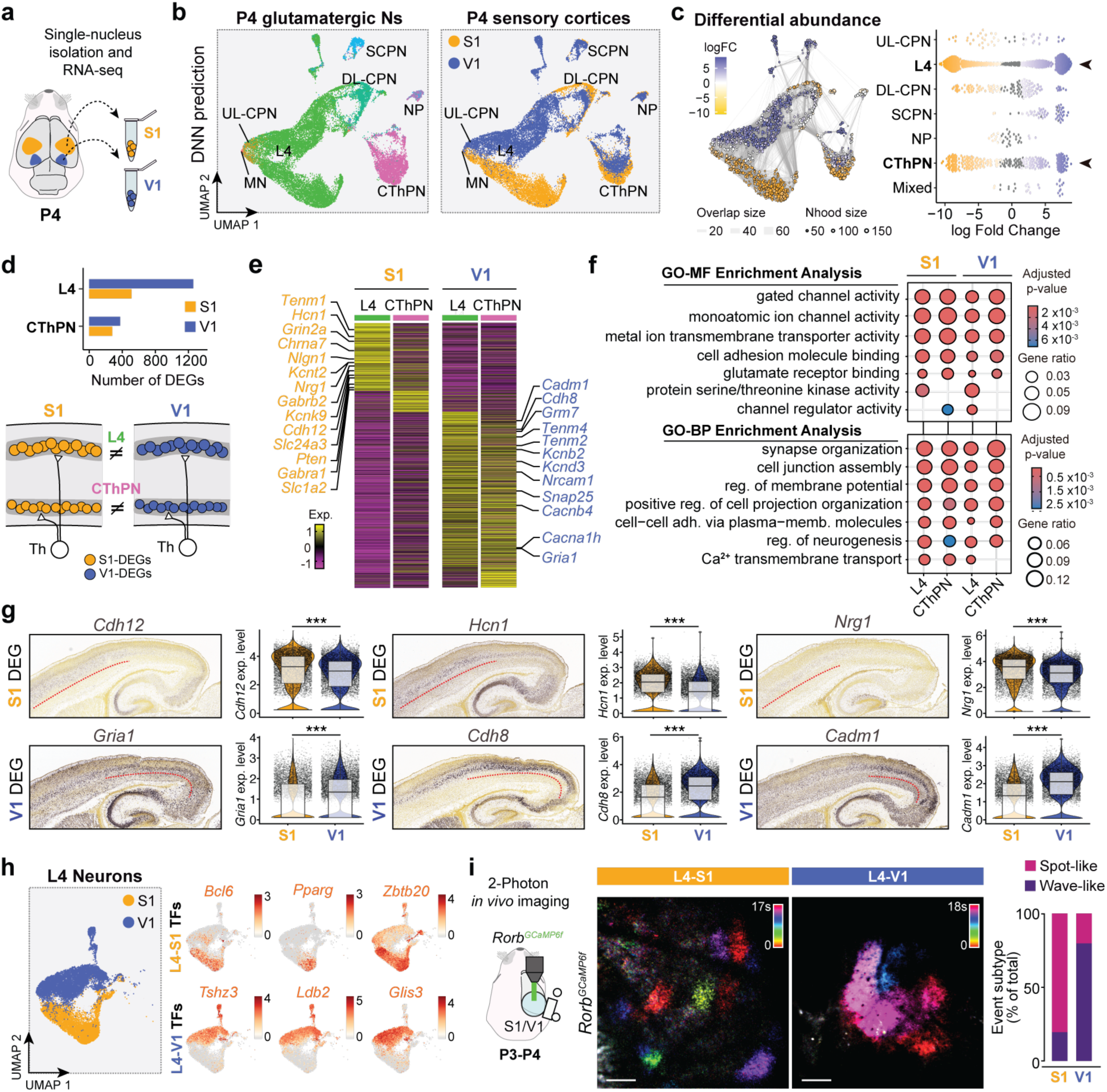
Cortex modality-specific transcriptomic differences lie in thalamo-recipient layers. **a,** Schematic of the experimental design. **b,** UMAP of glutamatergic neurons isolated from the P4 snRNA-seq dataset, colour-coded by deep neural network (DNN)-predicted cell type (left) and by cortical region (S1, orange; V1, blue; right). **c,** (Left) UMAP of P4 glutamatergic neuron neighbourhoods. Colours denote significant enrichment for S1 (orange) or V1 (blue). Point size reflects neighbourhood cell number; edge thickness indicates shared cells between neighbourhoods. (Right) Beeswarm plot showing neighbourhood enrichment for S1 and V1. Each point represents a neighbourhood (50–150 cells) with similar expression profiles. The x-axis denotes enrichment (log fold change). Colours indicate significance: grey, not significant; blue, V1-enriched; orange, S1-enriched. **d,** (Top) Number of differentially expressed genes (DEGs) in L4 and CThPN. (Bottom) Schematic summarizing main findings. **e,** Heatmap of normalized mean expression of region-specific DEGs in L4 and CThPN neurons. **f,** Gene ontology (GO) enrichment of S1 and V1 L4 genes, showing representative molecular functions (MF, top), and biological processes (BP, bottom). **g,** Sagittal *in situ* hybridization images illustrating regionally enriched DEGs in S1 and V1 at P4. Red dashed line marks the lower boundary of L4. Images from Allen Institute for Brain Science (https://developingmouse.brain-map.org). Violin plots show expression levels of L4 DEGs across S1 and V1. **h,** (Left) UMAP of L4 neuronal isolated population. (Right) Feature plots of representative transcription factors (TFs) differentially expressed between S1 and V1 L4 neurons. **i,** (Left) Schematic of the experimental design for L4-specific two-photon imaging. (Middle) Temporal colour-coded projection of spontaneous activity recorded in L4-S1 and L4-V1 neurons. (Right) Proportion of spot-like or wave-like events in L4 of S1 and V1. CThPN, corticothalamic projecting neurons; DL-CPN, deep-layer cortical projecting neurons; L4, layer 4 neurons; NP, near projecting neurons; Ns, neurons; P, postnatal day; S1, primary somatosensory cortex; SCPN, subcortical projecting neurons; UL-CPN, upper-layer cortical projecting neurons; UMAP, uniform manifold approximation and projection; V1, primary visual cortex. Scale bar 50 μm in **i**.

To determine where within the cortical circuit these differences arise, we assigned the isolated glutamatergic neurons to cortical subclasses using a pretrained deep-neural network (DNN) classifier^24^ (Fig. 1b). Differential abundance and expression analyses revealed that L4, the principal thalamo-recipient layer, contained most modality-specific genes, with a smaller contribution from L6 corticothalamic projecting neurons (CThPNs) (Fig. 1c). In S1, 512 genes were enriched in L4 and 277 in CThNs, whereas V1 L4 displayed 1,260 and 376 genes, respectively (Fig. 1d). These included regulators of synapse organization (e.g., *Grin2a* in S1; *Gria1* in V1)^25^ and cell-adhesion molecules (*Tenm1* and *Nlgn1*, in S1; *Cdh8*, *Tenm2* and *Tenm4* in V1)^26–32^ (Fig. 1e). Indeed, gene ontology (GO) analysis demonstrated that the most significantly enriched molecular functions and biological processes were related to neuronal activity, synapse organization, and cell adhesion (Fig. 1f; Fig. S2a,b). Pseudotime analysis confirmed comparable maturation in both cortices and therefore excluded that the transcriptomic differences reflect a delayed maturation of V1 relative to S1 (Fig. S2c-f). *In situ* hybridization and expression levels confirmed robust, layer-specific expression at P4 of representative markers, like *Hcn1* in S1 or *Cdh8* in V1 (Fig. 1g). Notably, nearly half of the 25 most differentially expressed genes (DEGs) encoded TFs or transcriptional regulators (e.g. *Bcl6*, *Pparg*, *Zbtb20* in S1; *Tshz3*, *Ldb2, Glis3* in V1 (Fig. 1h; Fig. S2g) with known roles in cortical development and neurodevelopmental disease^33–40^ (Table 1).

We next asked whether the transcriptomic differences in activity-related genes of excitatory neurons, such as those regulating synaptic signalling and membrane potential, are reflected in distinct activity profiles between S1 and V1. Wide-field calcium imaging in *Emx1;GCaMP6f* mice confirmed a functional correlate: S1 exhibited predominantly stationary, spot-like spontaneous activity events of <1 s, while V1 showed large propagating waves lasting 1–10+ s (Fig. S3). Two-photon calcium imaging restricted to L4 (*Rorb;GCaMP6f* mice^41^) recapitulated these differences (Fig. 1i), revealing L4 as both the molecular and functional locus of early modality identity. Together, these results demonstrate that S1 and V1 differ primarily in the transcriptional and activity signatures of their thalamo-recipient layers, highlighting L4 as a key locus for the emergence of cortical modality identity.

## Patterned thalamic activity refines, but does not establish, modality specific cortical identity

Given the concentration of modality-specific genes in thalamo-recipient layers, we asked whether their expression depends on developmental patterned thalamic activity. To test this, we used *Gbx2CreERT2;R26Kir2.1* (*Th^Kir^*mice hereafter) mice, in which thalamic relay neurons overexpress the inward rectifier potassium channel Kir2.1, altering thalamic activity^13–15^, and performed bulk RNA-seq and snRNA-seq on isolated S1 and V1 cortices at P4. Deconvolution of bulk RNA-seq profiles using cell-type signatures derived from the snRNA-seq data revealed that L4 was the layer most affected at the transcriptional level by the loss of thalamic synchronous activity, as further confirmed by the single-nucleus data (Fig. S4). We thus sought to investigate whether such changes impacted modality-specific transcriptional identity. Stratification by area of origin and differential abundance analysis revealed that S1 and V1 remain transcriptionally distinguishable in *Th^Kir^*mice, with major differences again localized to L4 and CThPN neurons (Fig. 2a,b). Nonetheless, a subset of modality-specific genes was lost in the absence of patterned thalamic activity: 15.2% (78/512) of S1 L4 genes and 37.8% (477/1260) of V1 L4 genes (Fig. 2c), demonstrating a direct but modest link between patterned thalamic activity and sensory-modality transcriptional identity.

**Fig. 2.**
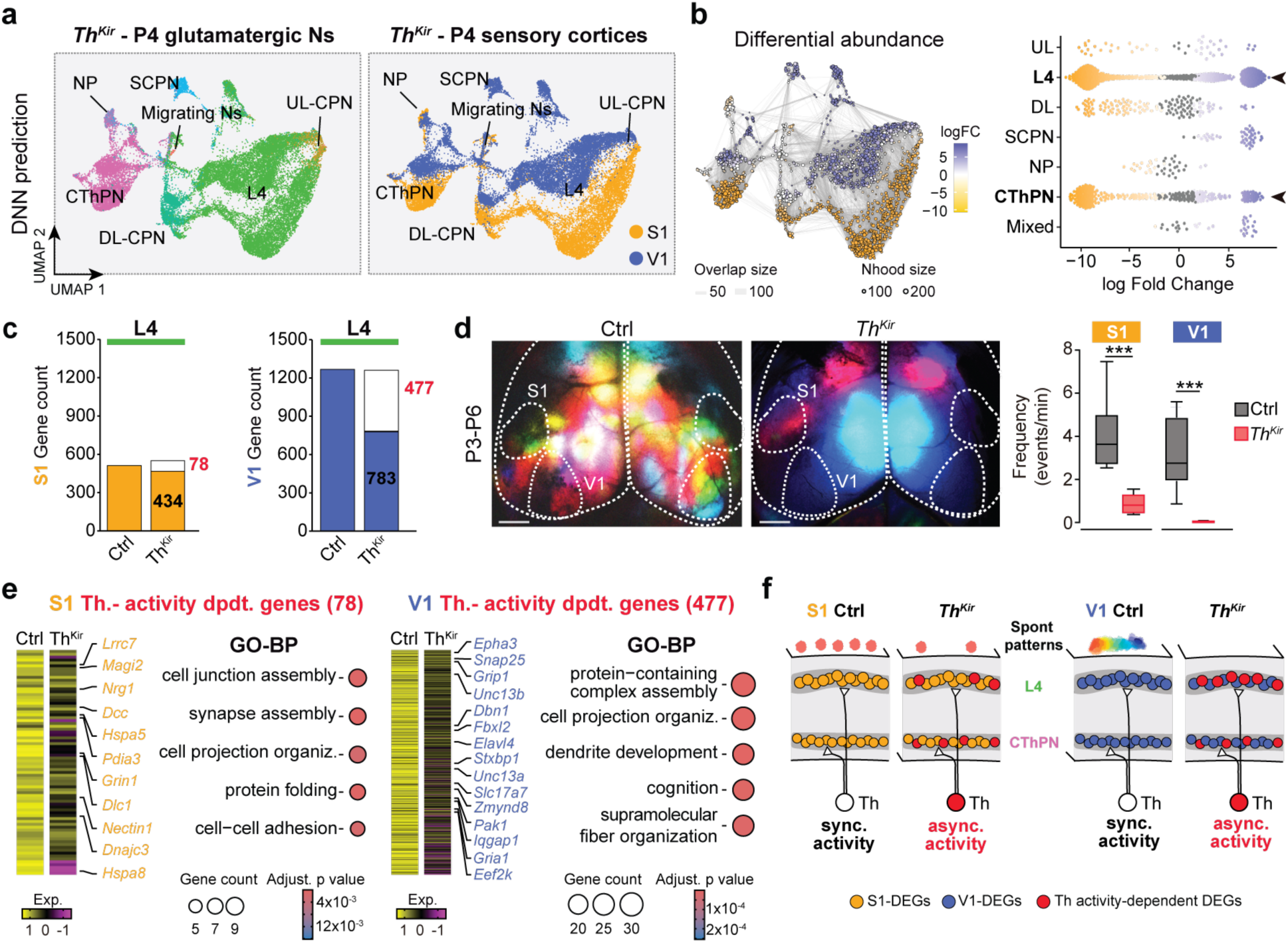
A subset of S1 and V1 identity genes is modulated by patterned thalamic activity. **a,** UMAP of the P4 snRNA-seq dataset from thalamic-Kir (*Th^Kir^*) mice, colour-coded by deep neural network (DNN)-predicted cell type (left) and by cortical region (S1, orange; V1, blue; right). **b,** (Left) UMAP of *Th^Kir^* glutamatergic neuron neighbourhoods. Colours denote significant enrichment for S1 (orange) or V1 (blue). Point size reflects neighbourhood cell number; edge thickness indicates the number of shared cells between neighbourhoods. (Right) Beeswarm plot showing neighbourhood enrichment for S1 and V1. Each point represents a neighbourhood (100–200 cells) with similar expression profiles. The x-axis denotes enrichment (log fold change). Colours indicate significance: grey, not significant; blue, V1-enriched; yellow, S1-enriched. **c,** Number of differentially expressed genes (DEGs) in L4 of S1 and V1 in control versus *Th^Kir^* mice. The number of thalamic activity-dependent DEGs (absent in *Th^Kir^*) is shown in red. **d,** (Left) Temporal colour-coded projection of a one-minute recording of spontaneous cortical activity in control and *Th^Kir^* mice at P4. (Right) Quantification of activity frequency in S1 and V1 from control (*n* = 10) and *Th^Kir^* (*n* = 6) mice at P4. **e,** Heatmaps of normalized mean expression and gene ontology biological processes (GO-BP) enrichment analysis for region-specific L4-S1 (left) and L4-V1 (right) activity-dependent DEGs affected in *Th^Kir^* mice. **f,** Schematic summary of the main findings. CThPN, corticothalamic projecting neurons; DL-CPN, deep-layer cortical projecting neurons; L4, layer 4 neurons; NP, near projecting neurons; Ns, neurons; P, postnatal day; S1, primary somatosensory cortex; SCPN, subcortical projecting neurons; UL-CPN, upper-layer cortical projecting neurons; UMAP, uniform manifold approximation and projection; V1, primary visual cortex. Scale bar 1000 μm in **d**.

Functionally, *Th^Kir^* mice exhibited a dramatic reduction in cortical spontaneous events and a near absence of propagating waves in V1 (Fig. 2d). Consistent with our previous findings of disrupted somatotopy in *Th^Kir^* mice^14^, retinotopic organization of geniculo-cortical projections was also perturbed, as revealed by DiI/DiD tracing in dLG thalamic nucleus (Fig. S5). Notably, the activity-dependent identity genes were enriched for processes related to cell projection organization (like *Nrg1* in S1 or *Epha3* in V1) and synapse and cell-junction assembly (like *Grin1* in S1 or *Snap25* and *Gria1* in V1) (Fig. 2e), supporting impaired circuit organization and maturation. These findings show that although transcriptional identity of S1 and V1 is largely preserved in the absence of patterned thalamic input, a subset of modality-specific genes is activity-dependent, and that proper cortical spontaneous activity and geniculo-cortical topography require a specific pattern of thalamic spontaneous activity (Fig. 2f).

## An embryonic, activity-independent cortical identity program predates thalamic innervation

Since most L4 identity genes persist in *Th^Kir^* mice, we hypothesized that modality identity is initiated intrinsically in the cortex before thalamic inputs target L4 neurons. At embryonic day 18 (E18), thalamic axons contact the subplate but have not yet reached L4^42,43^ (Fig. 3a; Fig. S6a-c), and S1 and V1 exhibit highly correlated spontaneous activity (Fig. 3b–d), indicating functional immaturity^44^. Nevertheless, snRNA-seq revealed incipient transcriptional divergence between S1 and V1, again concentrated in presumptive L4 and CThPN neurons (Fig. 3e-g; Fig. S6d-g). Remarkably, approximately half of these embryonic L4 DEGs persisted into P4 and, almost all were unaffected in *Th^Kir^* mice (Fig. 3h). We termed these time-stable, thalamic activity-independent cortical genes *protogenes*.

**Fig. 3.**
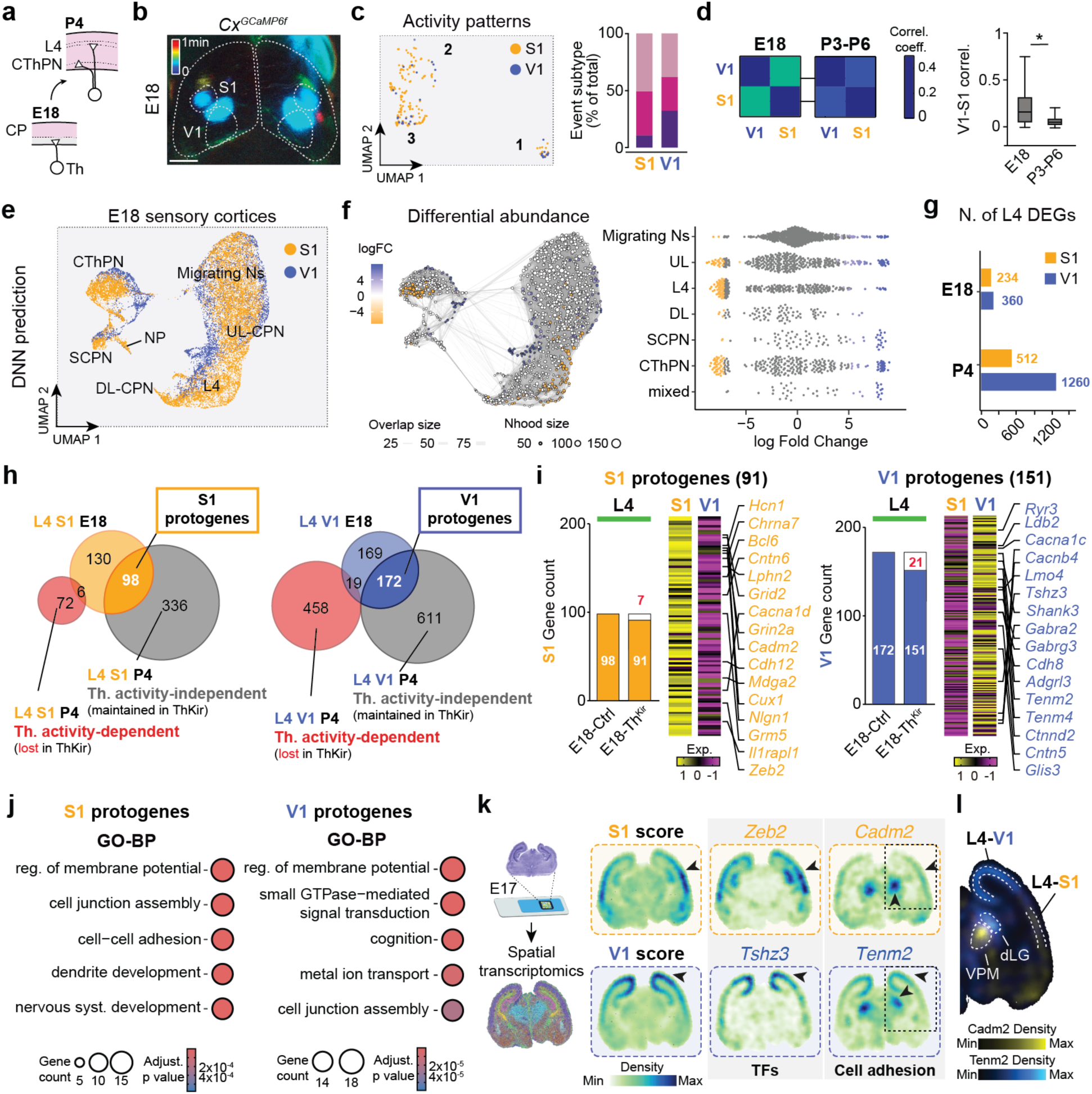
Embryonic functional coupling of S1 and V1 and emergence of modality-specific protogenes. **a,** Schematic illustrating the distinct influence of thalamic input on the cortex at E18 versus P4. **b,** Temporal colour-coded projection of one minute of spontaneous cortical activity recorded at E18. **c,** (Left) UMAP of calcium events detected in S1 (orange) and V1 (blue) at E18. (Right) Proportion of events belonging to each activity profile in S1 and V1. **d,** (Left) Developmental progression of S1-V1 activity correlations at E18 and P3-P6. (Right) Quantification. E18 (*n* = 16); P3-P6 (*n* = 10). **e,** UMAP of the E18 snRNA-seq dataset, colour-coded by cortical regions (S1, V1) and annotated with deep neural network (DNN)-predicted cell type. **f**, (Left) UMAP of E18 cortical cell neighbourhoods. Colours denote significant enrichment for S1 (orange) or V1 (blue). Point size reflects neighbourhood cell number; edge thickness indicates shared cells between neighbourhoods. (Right) Beeswarm plot showing neighbourhood enrichment for S1and V1. Each point represents a neighbourhood (50–150 cells) with similar expression profiles. The x-axis denotes enrichment (log fold change). Colours indicate significance: grey, not significant; blue, V1-enriched; orange, S1-enriched. **g,** Number of L4 differentially expressed genes (DEGs) between S1 and V1 at E18 and P4. **h,** Venn diagrams of S1-specific (left, orange) and V1-specific (right, blue) L4 DEGs at E18 and their overlap with P4 L4 DEGs that are thalamic activity-dependent (altered in *Th^Kir^*, red) or activity-independent (grey). **i,** Bar plots showing the number of S1 and V1 L4 DEGs at E18 in control and *Th^Kir^* mice (left). Activity-dependent DEGs (lost in *Th^Kir^*) are shown in red. Heatmaps showing normalized expression of S1 and V1 protogenes (right). **j,** Gene ontology (GO) enrichment of region-specific S1 and V1 L4 protogenes, showing representative biological processes (BP). **k,** (Left) Schematic of the E17 spatial transcriptomic analysis. (Right) Spatial density maps for S1 and V1 gene module scores, selected transcription factors and cell-adhesion molecules. **l,** Overlay of *Cadm2* and *Tenm2* spatial expression densities showing complementary distributions aligned with VPM-S1 and dLG-V1 territories. Arrowheads indicate regions of strong expression. CThPN, corticothalamic projecting neurons; DL-CPN, deep-layer cortical projecting neurons; L4, layer 4 neurons; NP, near projecting neurons; Ns, neurons; P, postnatal day; S1, primary somatosensory cortex; SCPN, subcortical projecting neurons; TFs, transcription factors; UL-CPN, upper-layer cortical projecting neurons; UMAP, uniform manifold approximation and projection; V1, primary visual cortex. Scale bar, 1000 μm in **b**.

Although at E18 there is no direct apposition of thalamic axons with L4 neurons it is possible that thalamic activity could exert its influence in gene expression through other mechanisms, like the activation of the subplate network^45^. However, only a small fraction of protogenes was altered in *Th^Kir^* mice at E18, 7,1% (7/98) in S1 and 12,2% (21/172), confirming their independence from thalamic patterned activity (Fig. 3i; Fig. S7). GO analysis revealed that protogenes were enriched for membrane excitability and, critically, cell-adhesion molecules (Fig. 3j), including cadherins, teneurins, latrophilins and members of the Eph-ephrin family, all previously implicated in axon guidance and synapse formation^26–32^. Spatial transcriptomics at an earlier timepoint, E17, validated early region-specific expression of these adhesion genes, many already localized to L4 (Fig. 3k). Strikingly, several protogenes showed matched expression between modality-specific thalamic nuclei and their corresponding cortical targets, for example, *Cadm2* in VPM and S1, and *Tenm2* in dLG and V1 (Fig. 3l). This matching pattern suggests that cell-adhesion codes defined by protogenes are likely to mediate modality-specific thalamocortical targeting.

## A combinatorial adhesion code mediates modality-specific thalamocortical partner matching

To test whether thalamic nuclei of the two sensory modalities exhibit adhesion signatures matching those of their cortical counterparts, we integrated our P4 cortical snRNA-seq dataset with a P2 thalamic atlas^46^ (Fig 4a; Fig. S8a-d). CellChat analysis^47,48^ revealed that L4 neurons had the strongest predicted ligand–receptor interactions with first-order thalamic nuclei, consistent with their role as primary thalamic targets (Fig. S8e). Among the cell-adhesion pathways mediating L4-thalamus interactions, the ADGRL family^49,50^ exhibited the highest interaction strength (Fig. 4b) and showed strong specificity with VPM and dLG neurons when compared to cortical neurons (Fig. 4c).

**Fig. 4.**
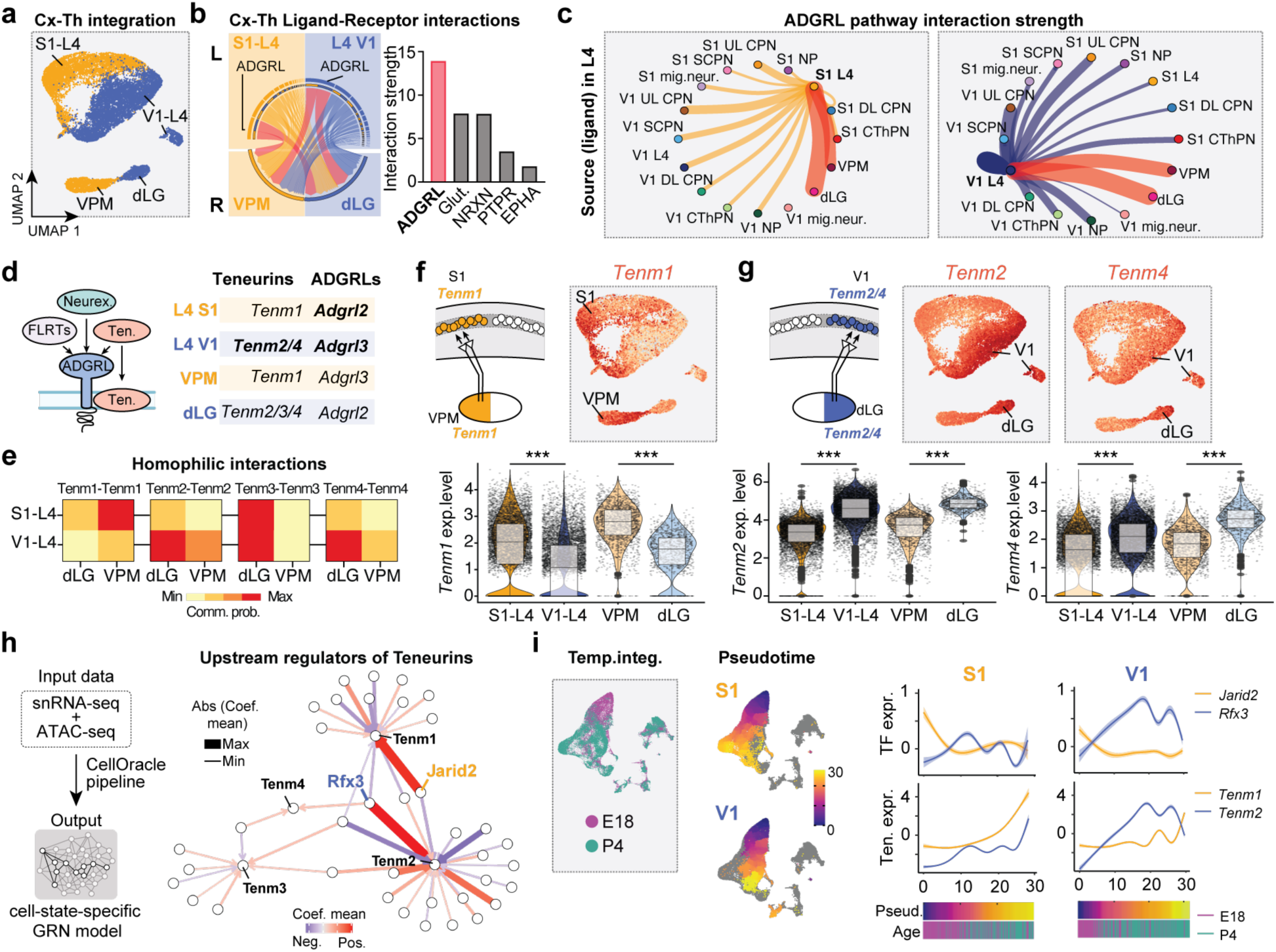
S1 and V1 protogene cell-adhesion signatures correspond with modality-specific thalamocortical signatures and innervation patterns. **a,** UMAP of the integrated P4 L4-cortex and P2 thalamus snRNA-seq datasets. **b,** (Left) Chord diagram showing major inter-areal pathways from L4 to thalamic neurons. Line width represents the communication probability. Outer bar colours are consistent with the signalling source and inner thinner bar colours represent the target that receives the signal from the corresponding outer bar. The outer bar length is proportional to the signal strength sent by the source. The inner bar length is proportional the signal strength received by the targets. Red lines mark the predominant pathway for each thalamus-cortex pair. (Right) Bar plot of the top 5 significant signalling pathways between thalamic and cortical L4 neurons. **c**, CellChat-inferred outgoing ADGRL pathway interactions from L4 neurons in S1(left) and V1 (right). Circle size indicates the number of cells per group; edge width and labels represent communication probability. Red lines mark the predominant cellular interactions. **d,** (Left) Diagram of the ADGRL family. (Right) Differentially expressed ADGRL-pathway members in L4 and thalamus across sensory modalities. Protogenes are shown in bold. **e,** Heatmaps showing the calculated communication probabilities for homophilic teneurin interactions between cortical L4 (S1, V1) and thalamic (dLG, VPM) neurons. **f,** (Top, left) Schematic summary of the expression findings. (Top, right) Feature plot showing matching modality-specific *Tenm1* expression in S1 and VPM. (Bottom) Violin plots showing expression levels of *Tenm1* across S1, V1, VPM and dLG. **g**, (Top, left) Schematic summary of the expression findings. (Top, right) Feature plots showing matching modality-specific *Tenm2* and *Tenm4* expression in V1 and dLG. (Bottom) Violin plots showing expression levels of *Tenm2* and *Tenm4* across S1, V1, VPM and dLG. **h,** (Left) Schematic of the approach for identifying the gene regulatory network (GRN) in L4 cortical neurons. (Right) Chord diagram of predicted upstream regulators of teneurins derived from CellOracle analysis. Line width represents the scaled mean coefficient score. Transcription factors are colour-coded by area specificity (S1, orange; V1, blue). Positive (red) and negative (indigo) regulatory influences are indicated by coefficient polarity. **i,** (Left) UMAP of the temporally integrated E18 and P4 cortical datasets. (Middle) Pseudotime trajectory of isolated cortical areas. (Right) Scaled expression dynamics of transcription factors and teneurins along pseudotime. Gene expression patterns were modelled using generalized additive models. CThPN, corticothalamic projecting neurons; DL-CPN, deep-layer cortical projecting neurons; dLG, dorsolateral geniculate nucleus; L4, layer 4 neurons; NP, near projecting neurons; Ns, neurons; P, postnatal day; S1, primary somatosensory cortex; S1, primary somatosensory cortex; SCPN, subcortical projecting neurons; UL-CPN, upper-layer cortical projecting neurons; UMAP, uniform manifold approximation and projection; V1, primary visual cortex. VPM, ventral posteromedial nucleus.

The ADGRL pathway comprises FLRT, neurexin, latrophilin and teneurin molecules (Fig. 4d), which interact to form trans-synaptic complexes involved in axon guidance and synapse formation^51,52^. Teneurins, in particular, mediate attractive homophilic recognition and synaptic partner-matching in both *Drosophila*^30,31^ and mice^32^. We therefore asked whether these proteins could mediate modality-specific thalamus–cortex interactions. Three out of four teneurin genes (*Tenm1*, *Tenm2* and *Tenm4*) showed modality-specific, thalamus–cortex matched expression that was preserved in *Th^Kir^* mice (Fig. 4d), indicating that their early expression is largely activity-independent. CellChat analysis of potential homophilic interactions showed that for Tenm1, 2 and 4, the highest predicted interaction probability occurred between each cortical region (S1 or V1) and its corresponding thalamic nucleus (VPM or dLG) (Fig. 4e). This pattern is consistent with a model in which homophilic interactions may contribute to modality-specific thalamocortical targeting. Analysis of expression patterns further revealed that *Tenm1* was enriched in S1-L4 and VPM, whereas *Tenm2* and *Tenm4* were enriched in V1-L4 and dLG (Fig. 4f,g).

To investigate upstream regulatory mechanisms, we applied the CellOracle^53^ pipeline to infer gene regulatory networks (GRNs) controlling cell-adhesion-related protogenes (Fig. 4h). Two TFs were identified as the principal regulators: Jarid2, enriched in S1, and Rfx3, enriched in V1. Jarid2 was predicted to positively regulate *Tenm1* and several S1-enriched adhesion genes (e.g., *Nlgn1*, *Sorcs1*) while repressing V1-specific genes. Conversely, Rfx3 was predicted to activate *Tenm2*, *Epha6*, and additional V1-enriched genes while repressing S1-specific transcripts (Fig. 4h; Fig. S9a,b).

Consistently, integration of E18 and P4 datasets and pseudotime analysis revealed coordinated temporal dynamics between teneurins and their predicted upstream TFs. *Tenm1* expression broadly tracked with *Jarid2*, whereas *Tenm2* expression strongly followed *Rfx3* (Fig. 4i; Fig. S9c,d). These developmental trajectories support a model in which early transcriptional programs establish lineage-specific adhesion signatures that are subsequently maintained postnatally.

Together, these data reveal a combinatorial thalamocortical cell-adhesion code, established embryonically and maintained postnatally, in which matched expression patterns of teneurins and related adhesion molecules align first-order thalamic nuclei with their appropriate cortical targets. This adhesion landscape, likely regulated by distinct TFs in S1 (Jarid2) and V1 (Rfx3), may contribute to the emergence and consolidation of modality-specific thalamocortical circuits.

## Discussion

Our findings provide a spatio-temporal dissection of the intrinsic genetic and extrinsic activity-dependent contributors to sensory cortical arealization. We identify two complementary classes of cortical genes: a core set of protogenes that define sensory-modality identity independently of thalamic activity and a subset whose postnatal expression requires patterned thalamic activity. Both gene sets are concentrated in L4, the principal thalamo-recipient layer, suggesting a two-step model for modality specification: intrinsically expressed protogenes establish an early molecular template that primes modality-specific thalamocortical targeting, while patterned thalamic activity subsequently refines cortical transcriptional identity and functional output (Fig. S10).

Sensory modalities are known to emerge functionally intermingled in the embryo and, thus, a somatosensory stimulus in the periphery triggers a cortical response in both S1 and V1^44^. Intriguingly, even if concomitantly activated, S1 and V1 occupy non-overlapping regions of the cortex, with modality-specific thalamocortical innervation. This suggests that the mechanisms driving cortical region identity and modality-specific thalamocortical targeting are tightly linked and established prior to functional segregation. Indeed, we identify a set of cortical protogenes differentially expressed between S1 and V1 from embryonic stages that regulate cell adhesion and could account for modality-specific thalamocortical targeting. Remarkably, spatial transcriptomics and ligand–receptor analyses show that thalamic nuclei display matching adhesion profiles, suggesting a reciprocal molecular architecture consistent with mechanisms of thalamocortical partner matching. Among these, teneurins stand out as compelling candidates for modality-selective matching^54,55^, given their homophilic binding properties and roles in synaptic partner selection^30–32^. Our GRN analysis identifies Jarid2 and Rfx3 as the two main upstream regulators of these cell-adhesion signatures in S1 and V1, respectively, offering a molecular entry point into the establishment of region-specific connectivity.

Using *Th^Kir^* mice, which lack synchronous spontaneous activity^14^, we reveal that protogenes are thalamic activity-independent, demonstrating that subcortical activity is dispensable for the establishment of broad cortical identity. Nevertheless, we found that thalamic activity regulates the correct expression of a subset of modality-specific genes, the emergence of distinct cortical spontaneous activity patterns, and the correct geniculo-cortical topographic organization. Together, our results overcome the long-standing protomap–protocortex debate by supporting a hybrid model. Intrinsic transcriptional programs generate a molecular protomap that guides incoming thalamic axons toward appropriate cortical territories, while patterned thalamic activity fine-tunes this framework to consolidate functional modality identity. This dual mechanism may explain why essential aspects of thalamocortical targeting are preserved in models with disrupted cortical layering^56^ or even sensory deprivation^44^, underscoring the robustness of modality-specific wiring strategies. Whether manipulating protogene expression or cortical activity can alter thalamocortical targeting specificity or reconfigure cortical identity arises an important question for future work.

Notably, several protogenes and activity-dependent genes identified here have been previously associated with human neurodevelopmental and psychiatric disorders, including autism spectrum disorders (ASD), schizophrenia, and bipolar disorder (Table 1). For instance, *GRIN2A*, an S1 protogene, carries variants linked to schizophrenia^57,58^, *CACNA1C*, a V1 protogene, is associated with schizophrenia, bipolar disorder and ASD^59–61^; and *DLG2* (PSD93), a thalamic activity-dependent S1 gene, has been repeatedly implicated in schizophrenia^62–65^. Notably, haploinsufficiency of *JARID2* or *RFX3,* the two TFs identified here as the principal regulators of modality-specific adhesion programs, results in neurodevelopmental disorders^66,67^. These observations raise the possibility that disruptions in early modality-specific molecular programs, and in their activity-dependent refinement, may contribute to disease susceptibility. This perspective highlights the value of considering early cortical arealization and thalamocortical signalling in understanding the origins of neuropsychiatric pathology.

**Table 1.** Identity genes among the Top 25 DEGs implicated in neural development and their clinical relevance. S1-specific genes are indicated in yellow, V1-specific genes are indicated in blue.

| Category | Gene | Neuro-developmental Role | Clinical Relevance | Identity gene subtype |
| --- | --- | --- | --- | --- |
| **Transcription Factors (DNA-binding)** | **ZBTB20** | Hippocampal neuron specification<sup>21</sup> | Developmental delay and neurodevelopmental disorders<sup>28</sup> |  |
|  | **BCL6** | Cortical development, and neurogenesis<sup>12,23</sup> |  | **Protogene** |
|  | **TSHZ3** | Corticostriatal development<sup>22</sup> | Autism spectrum disorder<sup>22,29</sup> | **Protogene** |
|  | **TOX** | Regulation of cortical neurogenesis and neurite outgrowth<sup>24</sup> |  |  |
| **Transcriptional Cofactors / Regulators (Non-DNA-binding)** | **LDB2** | L5 pyramidal neuron development<sup>25</sup> | Schizophrenia<sup>30</sup> | **Protogene** |
| **Synaptic Function, Vesicle Transport & Neurotransmission** | **SLC18A2** | Encodes vesicular monoamine transporter 2 (VMAT2) | Schizophrenia and mood disorders<sup>31</sup> |  |
|  | **HRH1** | Encodes Histamine receptor 1 | Schizophrenia<sup>32</sup> |  |
| **Cell adhesion** | **SORCS1** | Regulates neurexin and AMPA receptors surface trafficking<sup>26</sup> | Attention Deficit Hyperactivity Disorder (ADHD)<sup>33</sup> | **Protogene** |
|  | **CDH4** | Promotes pioneer axon outgrowth in ventral thalamus<sup>27</sup> |  | **Protogene** |

## Methods

### Animals

All transgenic lines in this study were maintained on an ICR/CD-1 background and genotyped by PCR. The day of the vaginal plug was stipulated as E0.5. The TCA-GFP *Tg* mouse, in which thalamocortical axons are labelled with GFP, was used to visualize primary sensory areas^20^. The Cre-dependent mouse line *R26^GCaMP6f^* was purchased from Jackson Laboratories (Stock number 024105) to conditionally express the calcium indicator by specific neuronal populations.

To express GCaMP6f in cortical glutamatergic neurons, *R26^GCaMP6f^* mice were crossed with *Emx1^Cre/+^* transgenic mice (*Cx^GCaMP6f^* mouse) from Jackson Laboratories (Stock number 005628). To express GCaMP6f specifically in layer 4 neurons, *R26^GCaMP6f^* mice were crossed with *Rorb^Cre/+^*transgenic mice (*Rorb^GCaMP6f^* mouse). The *Rorb^GCaMP6f^*mouse line was kindly provided by Dr. Marta Nieto and was originally obtained from Jackson Laboratories (Stock number 023526).

To desynchronize embryonic thalamic activity, the *R26^Kir2.1-mCherry 13^* was crossed with an inducible CreERT2 mouse line driven by Gbx2, an early specific thalamic promoter (Gbx2*^CreERT2/+^*)^68^. Double mutants are referred as *Th^Kir^* throughout the manuscript. To visualize cortical activity in the *Th^Kir^* condition we generated a triple mutant by crossing the *Th^Kir^* mouse with the SNAP25*^GCaMP6s^* mouse from Jackson Laboratories (Stock number 025111). To visualize the position of thalamic axons in the developing cortex, the tamoxifen-inducible Gbx2*^CreERT2/+^* mouse line was crossed with a R26tdTomato Cre-dependent mouse line from Jackson Laboratories (Stock number 007908). The resulting double mutants are referred in the manuscript as *Th^tdTomato^* mice.

Tamoxifen induction of Cre recombinase in the double/triple mutant embryos was performed by gavage administration of tamoxifen (5 mg dissolved in corn oil, Sigma) at E10.5 to specifically target all primary sensory thalamic nuclei. Tamoxifen administration in pregnant mice produces non-desirable side effects such as delivery problems and decrease survival of newborn pups^69^. To increase the survival rate of young pups, we administered 125 mg/Kg of progesterone (DEPO-PROGEVERA®) intraperitoneally at E14.5 and implemented C-section procedure at E19.5 (e18.5 in the case of *Th^tdTomato^* experiments). Pups were then placed with a foster mother. In all cases, the CreERT2-negative littermates were used as controls of the experimental condition.

All animal procedures were approved by the Committee on Animal Research at the Universidad Miguel Hernández and carried out in compliance with Spanish and European regulations.

### Tissue preparation and cellular dissociation for snRNA-seq experiment

To collect tissue from the cortices of animals at E18 and P4, animals were decapitated, and their brains were dissected out in RNase-free conditions to prevent RNA degradation. The brains (three brains were pooled for each sample) were collected in ice-cold artificial cerebrospinal fluid (aCSF) solution (119 mM NaCl, 5 mM KCl, 1.3 mM MgSO4, 2.4 mM CaCl2, 1 mM NaH2PO4, 26 mM Na2HCO3, and 11 mM glucose) and cut in the vibratome in 300 μm slices.

Single-nucleus isolation for the snRNA-seq experiment was performed based on the Demonstrated protocol for Nuclei isolation from tissue from 10xGenomics (CG000124). The frozen tissue were then suspended with 500 μL of 0.1× lysis buffer [10 mM Tris-HCl pH 7.4, 10 mM NaCl, 3 mM MgCl2, 0,2 U/μL RNase inhibitor and 0.1% IGEPAL CA-630 (i8896 Sigma-Aldrich)] for 5 min on ice, which was stopped with chilled 500 μL wash buffer [1% PBS without Ca/Mg+2 supplemented with 1% BSA and 0,2 U/μL RNase inhibitor] and the samples were pipetted 5 times using the P1000. The nuclei were then collected by centrifugation at 500× g for 5 min at 4 °C, rinsed with 200 μL wash buffer three times, and re-suspended in wash buffer. Isolated single nuclei were filtered using a cell strainer (35-μm pore size) and inspected under a microscope to ensure they were successfully dissociated into single cells for subsequent sequencing. Hoechst was added to label nuclei for FANS gating, using a FACSAria III cell sorter (BD Biosciences). 43.2 μl of nuclei were sorted directly into empty 1.5 ml Eppendorf for downstream processing on the 10x Genomics Chromium platform.

### snRNA-seq data generation

The single-nuclei RNA-seq libraries were constructed by following the 10x Genomics Chromium Next GEM Single Cell 3′ Reagent Kit v3.1 (Dual Index) following the manufacturer’s protocol (document number CG000315, 10x Genomics).

Uniquely barcoded RNA transcripts were reverse-transcribed. 3′ Gene Expression libraries were generated according to the manufacturer’s user guide with use of Chromium Next GEM Single Cell 3’ Kit v3.1, 4 rxns v3.1 kit (PN-1000269), Chromium Next GEM Chip G Single Cell Kit, 16 rxns (PN-1000127) and Dual Index Kit TT Set A (PN-1000215) (10x Genomics). 3′ Gene Expression Libraries was assessed using the High Sensitivity DNA Kit (#5067-4626, Agilent) on a Bioanalyzer 2100 (Agilent Technologies).

Transcriptome libraries were sequenced on an Illumina NovaSeq X platform at Novogene Co. Ltd. Genomics core facility (Munich, Germany), with 28-10-10-90 bp configuration for RNA-seq libraries.

### snRNA-seq data processing

Each dataset (E18 V1, E18 S1, P4 V1, and P4 S1) was sequenced with two biological replicates per condition. Single-nucleus RNA-seq data were processed using CellRanger (v8.0.1, 10x Genomics). Raw FASTQ files were aligned to the GRCm38 mouse genome reference with default parameters, and filtered gene-barcode matrices were imported into Seurat (v5.2.1) in R (v4.4.2) for downstream analysis.

Quality control was applied to retain nuclei with nFeature_RNA > 300 and mitochondrial content < 5%. Gene expression counts were normalized using the “LogNormalize” method. Dimensionality reduction was performed by principal component analysis (PCA) on the top 3,000 highly variable genes, and uniform manifold approximation and projection (UMAP) embeddings were computed from the first 30 principal components. Cell clustering was carried out using FindNeighbors and FindClusters with a resolution parameter of 0.3. Doublets were detected using a consensus approach with the scDblFinder (v1.12) and scds (v1.14) packages and excluded from downstream analyses. Biological replicates for each condition were then merged to generate the final datasets used for subsequent analyses.

E18 and P4 snRNA-seq datasets were normalized using SCTransform (*SCTransform* function, Seurat v5.2.1). Integration was performed with the Harmony algorithm (*HarmonyIntegration*), after PCA on the top 3,000 highly variable genes. Harmony-corrected embeddings were used to construct a shared nearest-neighbour graph and to compute dimensional projections with 30 dimensions and the first 30 principal components.

To identify marker genes distinguishing clusters, we performed differential expression analysis using Seurat’s FindAllMarkers function on the RNA assay, applying a log2 fold change threshold of 0.25 and a minimum fraction of expressing cells of 0.1, with p-values adjusted using the Bonferroni correction. Cell clusters were annotated using canonical markers for neurons, astrocytes, oligodendrocyte precursor cells (OPCs), and microglia. Furthermore, for differential expression analysis between conditions (Control versus *Th^Kir^*) and neuronal layer cell types, we applied a simultaneous statistical significance threshold (Benjamini-Hochberg (BH) adjusted P-value < 0.1), absolute log2 fold change (log2FC) > 0.322 and min.pct > 0.1.

### Cell type abundance and neighbourhood changes between conditions

The miloR R package (v.1.6.0)^70^ was used to assess differential abundance within overlapping cell neighbourhoods, ranging in size from 50–200 cells each. For each cortical area (S1 and V1), the expression matrix was subset to include only the top 3,000 most variable genes within cells from areas, PCA was performed on that submatrix and the result was passed into a MiloR object for further analysis. miloR’s buildGraph function was run with k = 30 and d = 30, followed by makeNhoods and calcNhoodDistance. Differential abundance testing was performed for V1 versus S1 cells within each neighbourhood using testNhoods with a design of ∼Area. Significance is reported as a spatial FDR value calculated by miloR based on differential abundance P-values and the relationships between neighbourhoods.

### Gene ontology over representation analysis

Gene ontology (GO) over-representation analysis was performed for biological processes and molecular functions using the enrichGO function from the clusterProfiler package (v4.6.2), with the following parameters: pAdjustMethod = “BH”, qvalueCutoff = 0.01, minGSSize = 10 and maxGSSize = 500.

### Deep neural network prediction

Cortical cell identities were predicted using a feedforward neural network trained on a temporal single-cell transcriptomic dataset^24^ collected from the S1 cortical area during development (E10–P4) and generated with 10x Genomics technology. The dataset was downsampled to 2,000 cells per cell-type/identity class and split into training (80%) and test (20%) subsets. The sequential model was implemented and trained using the *Keras* (v2.15) and *TensorFlow* (v2.16) R packages in a multi-class categorical framework. The model was trained to classify cells into the annotated classes and layers of the original dataset, including glutamatergic neurons, GABAergic neurons, cycling glial cells, intermediate progenitors, astrocytes, oligodendrocytes, microglia, and pericytes. The glutamatergic neuron class was further subdivided into migrating neurons, upper-layer cortical projection neurons, layer 4 neurons, deep-layer neurons, subcortical projection neurons, corticothalamic projection neurons, and projecting neurons. Model evaluation on the test set yielded an accuracy of 97.41%, indicating robust prediction performance.

### Pseudotime trajectory analysis

Pseudotime trajectories were constructed using the Monocle3 workflow (v1.3.7). The merged/integrated Seurat object was converted into a cell_data_set using the SeuratWrappers package (v0.3.5). Dimensionality reduction and cell clustering was based on the UMAP embedding from the Seurat object. The principal graph was learned with learn_graph function (use_partition = FALSE), and pseudotime was assigned using the order_cells function with default settings. Scaled gene expression dynamics along pseudotime were modelled with generalized additive models (GAMs) using the formula *Scaled expression ∼ s(pseudotime, bs=”cs”, k=7)*, where *s* denotes a smooth spline function. Smoothed expression curves were used to visualize temporal trajectories of selected genes.

### Microdissection and RNA isolation for bulk RNA-seq

To capture gene expression changes at high-throughput genome wide level, we collected tissue from the S1 and V1 cortical areas of P4 Ctrl and *Th^Kir^*samples. Animals were decapitated, and their brains were dissected out in RNase-free conditions to prevent RNA degradation. The brains (four brains were pooled for each sample) were collected in ice-cold artificial cerebrospinal fluid (aCSF) solution (119 mM NaCl, 5 mM KCl, 1.3 mM MgSO4, 2.4 mM CaCl2, 1 mM NaH2PO4, 26 mM Na2HCO3, and 11 mM glucose) and cut in the vibratome in 300 μm slices. The S1 and V1 areas were rapidly microdissected under a stereo microscope. The bulk tissue was immediately transferred to lysis buffer of the Rneasy® Micro Kit (Qiagen, 74004) for total RNA extraction, following the manufacturer’s instructions. RNA quality was measured for all samples using an Agilent Bioanalyzer 2100 system, and only samples with RNA Integrity Number (RIN) > 8 were used for library construction.

### Library preparation and bulk RNA-seq

Library construction and sequencing were performed at Novogene Co. Ltd. Genomics core facility (Munich, Germany). cDNA multiplex libraries were prepared using a custom Novogene NGS RNA Library Prep Set (PT042) kit. Briefly, mRNA was purified from total RNA using poly-T oligo-attached magnetic beads. After fragmentation, the first strand cDNA was synthesized using random hexamer primers followed by the second strand cDNA synthesis. The library was ready after end repair, A-tailing, adapter ligation, size selection, amplification, and purification.

The library was checked with Qubit and real-time PCR for quantification and bioanalyzer for size distribution detection. Libraries were pooled and sequenced in 2x150bp paired-end mode on a S4 flowcell in the Illumina Novaseq6000 platform. A minimum of 40 million reads were generated from each library.

### Bioinformatic analysis of the bulk RNA-seq

RNA-seq analysis were performed as previously described^,44,71^ with minor modifications: quality control of the raw data was performed with FastQC (v.0.11.9). RNA-seq reads were mapped to the Mouse genome (GRCm39) using STAR (v2.7.9a)^72^ and SAM/BAM files were further processed using SAMtools (v1.15). Aligned reads were counted and assigned to genes using Ensembl release 104 gene annotation and FeatureCounts, Subread (v2.0.1). Normalization of read counts and differential expression analyses were performed using DESeq2 (v1.32)^73^ Bioconductor (v3.15)^74^ in the R statistical computing and graphics platform (v4.2.2 “Innocent and Trusting”).

In the analysis of Controls and *Th^Kir^* datasets generated for this study, significantly Differentially Expressed Genes (DEGs) were identified using a simultaneous statistical significance threshold (Benjamini-Hochberg (BH) adjusted P-value < 0.05) and absolute log2 fold change (log2FC) > 0.322 by shrunken log2FC using the adaptive T prior Bayesian shrinkage estimator “apeglm” ^75^. Hierarchical clustering analysis was performed using “Euclidean” distance and “Complete” clustering methods metrics to visualize significantly upregulated and down-regulated genes.

### Module scoring for feature expression programs

DEGs lists derived from bulk RNA-seq analyses were used to compute module scores representing feature expression programs in single-cell data. Module scores were calculated with the Seurat AddModuleScore function, which assigns a score to each cell based on the average expression of the input gene set, relative to control gene sets matched for expression level and gene count. For each DEG list, module scores were computed across all cells in the dataset and subsequently used to assess cell-type–specific or condition-specific enrichment of bulk-derived transcriptional signatures.

### Spatial transcriptomic

Spatial transcriptomic profiling was performed using the Curio Seeker 10×10 Kit (Curio Biosciences, CB-000578-UG v1.3). Brains at E17.5 were washed in PBS with 2 U/mL RNase inhibitor, incubated in tissue-freezing medium (Electron Microscopy Sciences), embedded in BEEM flat molds, and snap-frozen on dry ice. Samples were cryosectioned at −18 °C using a Leica cryostat (CM1860 UV, Leyca Biosystems) at 10 μm thickness. Sections were carefully mounted on the Curio Seeker 10×10 mm tile and melted by placing a finger under the tile. A 30-μm-thick section of the CryoCube was placed on top of the tissue section and melted similarly.

RNA extraction and library preparation followed the manufacturer’s protocol. Tagmentation was performed using 4.8 ng cDNA and the Nextera XT Library Prep Kit (Illumina, FC-131-1024). Libraries were pooled and sequenced on a NovaSeq X Plus platform (Illumina) using paired-end 150 bp reads (150–8–8–150 bp configuration). Spatial gene expression matrices were processed using the Curio Seeker pipeline (v3.0).

### Cell-cell interaction analysis

Cell-cell communication analysis was performed in R using CellChat (v2.2.0)^47,48^ to identify ligand-receptor interactions between thalamic and cortical neurons. A publicly available P2 thalamic dataset previously generated in the lab^47^ was used to define neuronal clusters corresponding to the visual (dLG) and somatosensory (VPM) thalamic nuclei. Two integrated datasets were then constructed: (i) P4 S1 and V1 neurons combined with P2 dLG and VPM nuclei (24,975 nuclei), and (ii) P4 S1 L4 and V1 L4 neurons combined with P2 dLG and VPM nuclei (14,293 nuclei). Dimensionality reduction was performed by PCA on the top 2,000 highly variable genes, and UMAP embeddings were computed from the first 20 principal components. Cell–cell communication networks were inferred following the standard CellChat pipeline (https://github.com/jinworks/CellChat) and using the CellChatDB to which we manually added 4 ligand-receptor pairs based on literature. Global signalling interactions were visualized with the netVisual_circle function, while the netVisual_chord_gene function was used to display major ligand–receptor pathways mediating communication between thalamus and cortex. The pheatmap function was used to plot the communication probabilities of homophilic interactions between teneurins.

### Gene regulatory network (GRN) inference and visualization

GRNs were inferred using CellOracle (v0.17.2)^53^ following the recommended pipeline (https://morris-lab.github.io/CellOracle.documentation/). For computational efficiency, 3,000 cells and 3,000 highly variable genes were downsampled from the snRNA-seq dataset. Preprocessing and clustering were performed with Scanpy (v1.9.6), and diffusion maps, PAGA, and force-directed graphs were computed using default parameters.

A consensus ATAC-seq peak file generated by the CellOracle pipeline was processed to obtain a BED file indicating genomic coordinates of ATAC peaks. Peaks overlapping transcription start sites (TSS) were annotated. Using the CellOracle wrapper for gimmemotifs (v0.17.0), a base GRN was constructed by identifying TF-binding motifs within ±2 kb of all gene promoters. TF–target interactions were then filtered using a significance threshold of P-value < 0.001, with the absolute regression coefficient (“coef_abs”) serving as the weighting metric, and a maximum of 2,000 TF–target links retained.

For visualization, the inferred GRNs were imported into R and represented as graph objects using the igraph package (v2.0.3). Network layouts were computed with force-directed algorithms, and node/edge attributes (TF identity, target class, edge weight) were mapped onto visual properties. Final publication-quality network figures were generated using the ggraph package (v2.2.1), enabling visualization of global network topology and highlighting of TF-specific subnetworks.

### Assignment of cortical territories for mesoscale calcium imaging

The perimeter of the primary sensory cortex (S1) was determined using the cortical responses elicited by mechanical stimulation of five sites of the whisker pad, as described previously^14^. The perimeter of the primary visual cortex (V1) at E18 and P0 was predicted by scaling down and superimposing the limits of the sensory territories observed in the TCA-GFP transgenic line^20^ at P2 and using the responses to whisker pad stimulation as a reference for S1.

### *In vivo* mesoscale calcium imaging

At E18, embryos were retrieved from the uterus and kept at 35°C. E18 and P0 pups were immobilized with soft clay. Mice between P3 and P6 were anesthetized with ice and their scalp was removed surgically. A 3D-printed plastic holder was glued to the skull with cyanoacrylate adhesive and dental cement and affixed to a ball-joint holder to immobilize the head. Pups were placed on a controlled temperature heating pad to keep their body temperature at 32-34°C. We used a 16-bit CMOS camera (Hamamatsu ORCA-Flash 4.0) coupled to a stereo microscope (Stereo Discovery V8, Zeiss) that provided 470 nm LED illumination to record calcium activity. For *Cx^GCaMP6f^* mice, images were acquired with a frame size of 1024x1024 pixels using a macro magnification of 1.25x at E18-P0 and of 1x at P3-P6. Thus, the spatial resolution was 10.64 μm/pixel and 13.16 μm/pixel, respectively. Image frames were acquired continuously for at least 5-10 min at a rate of 3.33 frames per second (300 ms frame period). An average of 3 videos was acquired per animal.

### *In vivo* two-photon calcium imaging

To examine functional differences in spontaneous activity between S1 and V1, *Rorb^GCaMP6f^* pups with cranial windows over their corresponding cortical regions (S1 or V1) underwent two-photon calcium imaging at P3 or P4. Craniotomies^20,76^ were performed on the same day of imaging under isoflurane anaesthesia (2%). The skull over the target area was carefully removed without damaging the dura and replaced with a 3-mm round coverslip (Warner Instruments). Cortex buffer^77^ was applied throughout the surgery. A custom 3D-printed plastic holder was glued to the head using dental cement. Following surgery, pups were returned to their littermates on a 37°C heating pad for at least 1 h. For imaging, pups were head-fixed under the microscope without anaesthesia. Body temperature was maintained using a small heating pad, and pups were covered with a soft cotton blanket to minimize stress. Calcium signals were recorded using a Ti:sapphire laser (Axon 920-2 TPC, Coherent) and a GaAsP photomultiplier (Thorlabs) through a 16x immersion objective (Nikon). Images were acquired at a resolution of 1024x1024 pixels with a frame rate of 15.21 Hz (1x) or 15.23 Hz (2x), for a total of 2000 frames per recording. Each session lasted ∼2 minutes, with a total imaging time of ∼15 minutes per pup. Multiple imaging sessions were obtained from each animal to maximize data collection while minimizing animal use. No significant body weight loss was observed following either craniotomy or imaging.

### Analysis of *in vivo* spontaneous activity

Spontaneous activity in the cortex was analysed *in vivo* using custom scripts written in MATLAB™ (The MathWorks, Natick, MA) and Python. To delineate the perimeters of S1 and V1, the acquired videos were aligned with a reference map obtained from the cortex of TCA-GFP mice at P2 (for E18) or P6 (for P4–P5 brains). To facilitate alignment, the boundaries of the barrel field were identified by stimulating the whisker pad. Each video was preprocessed as follows: frames containing motion artifacts were removed and, when necessary, spatial downsampling was applied until reaching a final resolution of 512 × 512 pixels. ΔF/F₀ was calculated using the 10th percentile of the signal as F₀. Independent component analysis (ICA, MATLAB implementation) was then applied to remove residual artifacts (e.g., hemodynamic signals, unresolved motion artifacts), and the independent components were manually classified^78–81^. Baseline correction was performed with the built-in MATLAB function *msbackadj* (window size = 20 s, step size = 5 s), and a spatial median filter was applied (filter size [5,5]). Subsequently, images were segmented and converted to a binary scale using a global thresholding approach, where the threshold was defined as an estimate of background noise (median absolute deviation of fluorescence divided by 0.6745)^82^, multiplied by a fudge factor of ∼20. Segments related to the built-in function *bwconncomp* using default connectivity (26 for 3D arrays), such that overlapping areas in contiguous frames were considered part of the same calcium event. Finally, calcium event detection was refined manually: events with very small intensity, size, or duration were discarded; events outside the ROI were excluded; and merged events were resolved through a heuristic, supervised implementation of an ICA-based method^83^ applied to the minimovie containing the merged events.

We then analysed several features of the detected calcium events. For each event, we quantified parameters related to duration (event duration, width at 50% of the prominence, rise and decay times), shape (eccentricity, length of the major and minor axes, area of the maximum projection), signal intensity (prominence, baseline), and movement (displacement, defined as the distance between the event centroid at onset and at the end, and arc length, defined as the trajectory length of the centroid). Unless otherwise stated, data from the left and right hemispheres of the same animal were pooled. Finally, the active fraction of each brain region was computed as the percentage of active pixels within the region, and pairwise correlations between brain regions were calculated based on the active fraction.

### Clustering of spontaneous activity

A total of 12 features (see above) were used to characterize 649 events recorded in two cortical regions from 7 control mice at 2 developmental stages (3 control E18, 4 control P3-P6). For each calcium event, we extracted twelve features grouped in four modules—duration, shape, signal intensity and movement— and applied machine learning-based classification. These features were explored using histograms and descriptive statistics. As a preprocessing step, the distributions of all the features were transformed (Yeo-Johnson transformation) and scaled (standard scaling mean=0 and variance=1). The twelve dimensions of the dataset were reduced and visualized in Python using the UMAP algorithm^84^, applying the following parameters: number of neighbours equals 10, minimum distance equals 0.12 and a cosine distance metric. UMAP visualization suggested the existence of at least three different groups of events in the data set, identified by applying a hierarchical clustering algorithm to the pre-processed dataset (Ward linkage and Euclidean distance). As a result, each group was characterized by a vector (or profile) of twelve elements that contained a standardized version of each feature for all the events within the group. Three major activity profiles were identified in developing sensory cortex. Profile 1 corresponded to propagating wave-like events covering large cortical areas and lasting 1–10+ s, whereas profiles 2 and 3 represented smaller, stationary events of <1 s (profile 3 slightly longer).

### Dye-tracing studies

For axonal tracings at P15, animals were perfused with 4% PFA in PBS, and their brains dissected out and post-fixed overnight. Small DiI (1,1′-dioctadecyl 3,3,3′,3′-tetramethylindocarbocyanine perchlorate; Invitrogen) and DiD (1,1’-Dioctadecyl-3,3,3’,3’-Tetramethylindodicarbocyanine, 4-Chlorobenzenesulfonate Salt; Invitrogen) crystals were inserted under a stereo fluorescence microscope (MZ10 F, Leica) at the same distance from each other in either the elevation or azimuth orientation in V1 and allowed to diffuse for 4 weeks at 37 °C in PFA solution. Vibratome sections (80 μm thick) were obtained and counterstained with the fluorescent nuclear dye DAPI (Sigma-Aldrich). Five sections per animal were scanned in 3mm step-stacks using a Leica K5 camera in a Leica DMi8 microscope with a 10x objective.

### Immunofluorescence assay

For immunohistochemistry of the embryonic tissue, brains were dissected at E18 and immediately fixed in 4% PFA overnight. Brains were embedded in 3% agarose (in 0.01 M PBS) and sectioned at 80 μm thickness tangentially using a Leica VT1200S vibratome. Antigen retrieval was carried out by incubating the slices for 15 min at 80°C (300 rpm) in 10 mM sodium citrate buffer (0.05% Tween 20, pH 6). After that, the slices were cooled on bench top for 20 minutes, then washed in PBS and incubated for 1 h at room temperature in a blocking solution containing 1% bovine serum albumin (BSA) (Sigma) and 0.1% Triton X-100 (Sigma) in PBS. Afterwards, primary antibody incubation was performed overnight at 4°C with the following primary antibodies: rabbit anti-RFP (1:1000, Rockland, 600-401-379) and mouse anti-Satb2 (1:250, Abcam, ab51502). Sections were then rinsed in PBS and incubated for 2 h at room temperature with secondary antibodies: Alexa647 donkey anti-mouse (1:500, Invitrogen, A31571) and Alexa546 donkey anti-rabbit (1:500, Invitrogen, A10040). Nuclei were counterstained using the fluorescent nuclear dye 4′,6 diamidino-2-phenylindole (DAPI) (Sigma-Aldrich). After that, the sections were washed in PBS and mounted in Fluoromount-G (SouthernBiotech). Images were acquired with a Leica K5 camera in a Leica DMi8 microscope, with a 5x or 40x magnification.

### Fluorescence intensity analysis

To analyse the absence of TCAs in upper cortical layers at E18, fluorescence profiles of Satb2 and Gbx2 were measured using ImageJ. Cortical area S1 was localized with the help of the Allen Mouse Brain Atlas. Prior to analysis, pixel properties were adjusted to 0,324 μm/pixel. For the analysis of S1, the region of interest (ROI) was set at 550,55 μm in length, while for V1 the ROI was set at 460,47 μm. Using the “plot profile” option in ImageJ, a plot of the fluorescence intensity across the length of the ROI was generated.

### Statistics

Statistical analysis was carried out in GraphPad Prism 8, IBM SPSS Statistic 27 and Matlab. A Kolmogorov-Smirnov normality test was run for all datasets from activity and topography experiments. For independent data that followed a normal distribution, an unpaired two-tailed Student’s *t*-test was run to compare two groups. For independent data that did not follow a normal distribution, a Mann-Whitney *U*-Test two-tailed test or Wilcoxon rank sum test was used to compare two groups.

### Quantifications

Main Figures: In Fig. **1g**: Differential expression analysis between S1 and V1 areas in L4 neurons using a negative binomial test. *** adj. p<0.001. In Fig. **2e**: S1 Frequency, ctrl vs *Th^Kir^*, Mann-Whitney *U*-Test. \*\*\**P* < 0.001. V1 Frequency, ctrl vs *Th^Kir^*, Student’s *t*-test. \*\*\**P* < 0.001. In Fig. **3d**: V1-S1 correlation, E18 vs P3-P6, Student’s *t*-test. \**P* < 0.05. In Fig. **4f,g**: Teneurin expression levels across S1-L4, V1-L4, VPM and dLG, Kruskal-Wallis test and post hoc pairwise comparisons were conducted using Dunn’s test, with p-values adjusted using the Benjamini–Hochberg procedure. *** adj. p< 0.001 Supplementary Figures: In Fig. S**5b**: % dLG overlap in azimuth, ctrl vs *Th^Kir^* Mann-Whitney *U*-Test. \*\*\**P* < 0.001. % dLG overlap in elevation, ctrl vs *Th^Kir^* Student’s *t*-test. \*\*\**P* < 0.001.

## Data availability

All data supporting the findings of this study are available within the article and its Supplementary Information. The raw and processed snRNA-seq data from cortical areas at E18 and P4 stages, snRNA-seq data from thalamus at P2 stage, bulk RNA-seq data from cortical areas at P4 stage and the spatial transcriptomic data at E17 generated in this study have been deposited in the NCBI Gene Expression Omnibus and will be available upon publication. Additional data and materials necessary for reanalysis are available from the corresponding author upon request.

## Acknowledgments

We thank Belén Andrés Bayón, Luis Rodríguez Malmierca and Emily Wilson for technical support. We are also grateful to members of the López-Bendito laboratory for stimulating discussions throughout the course of this work. During the preparation of this work the authors used ChatGPT to streamline some parts of the text. After using this tool, the authors reviewed and edited the content as needed and take full responsibility for the content of the publication. This study was supported by: European Research Council under the European Union’s Horizon 2020 research and innovation programme (ERC-2021-ADG-101054313, SPONTSENSE) (G.L-B.); PID2021-127112NB-I00 and PID2024-155963NB-I00 grants (G.L-B.) from the MCIN/AEI /10.13039/501100011033; Generalitat Valenciana, Conselleria d’Educació, Universitats i Ocupació (PROMETEO 2021/052) (G.L-B.); T.G-V hold a Junior Research Fellowship from La Caixa Foundation (LCF/BQ/PR23/11980050) and at present a RYC2023-042550-I from the MCIN/AEI /10.13039/501100011033; L.P-A. hold a PRE2019-087666 from the MCIN/AEI /10.13039/501100011033; The Institute of Neurosciences is a Severo Ochoa Centre of Excellence (grant CEX2021-001165-S), funded by MCIU/AEI/10.13039/501100011033.

## Author contributions

Conceptualization, T.G-V, G.L-B; Methodology, T.G-V, L.P-A, L.W, D.V, D.T, M.A-M, I.F-J, FJ.M. Investigation, T.G-V, L.P-A, L.W, D.V, D.T, M.A-M, I.F-J, FJ.M. Visualization, T.G-V, L.P-A, L.W, D.V, D.T, M.A-M, I.F-J, FJ.M, G.L-B. Funding acquisition, G.L-B. Project administration, T.G-V, G.L-B. Supervision: T.G-V, G.L-B. Writing – original draft, L.P-A, T.G-V, G.L-B. Writing – review & editing, T.G-V, L.P-A, I. F-J, FJ.M, G.L-B.

## Supplementary Figures

**Figure S1.**
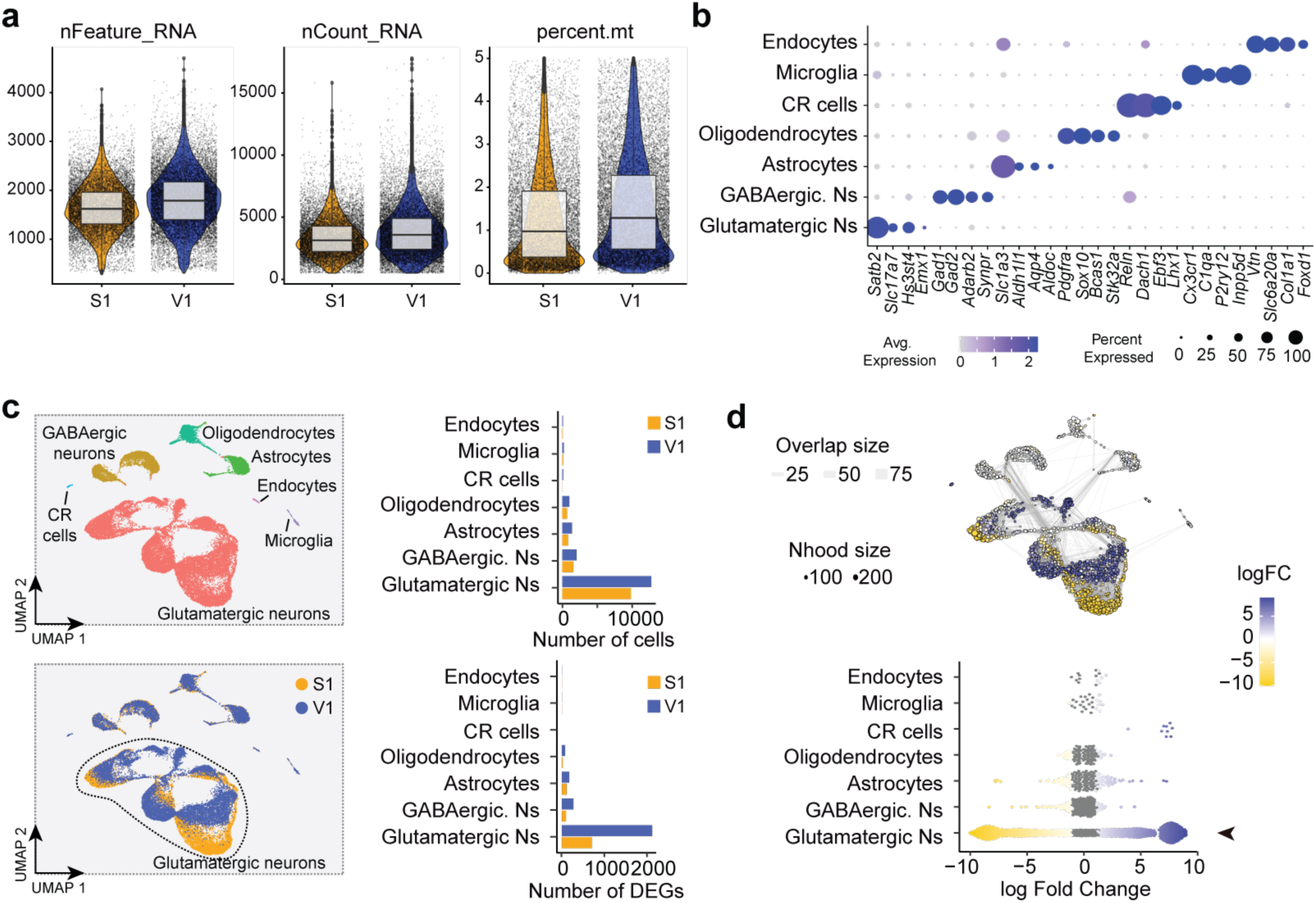
Cell type composition of S1 and V1 in the P4 snRNA-seq dataset. **a,** Data quality metrics of the control snRNA-seq libraries at postnatal day 4 (P4). Violin plots display distributions of total genes detected, total RNA reads, and percentage of mitochondrial gene counts (from left to right) split by cortical regions (S1, orange; V1, blue). **b,** Dot plot of representative marker genes defining major cortical cell classes. **c,** (Top) UMAP of the P4 snRNA-seq dataset colour-coded by identified cell populations (left). Bar plot showing the number of cells in each specific cortical region (S1, orange; V1, blue, right). (Bottom) UMAP of the P4 snRNA-seq dataset colour-coded by cortical region of origin (left). Bar plot showing the number of differentially expressed genes (DEGs) per cortical region (S1, orange; V1, blue, right). **d,** (Top) UMAP of cortical cell neighbourhoods at P4 (left). Colours denote significant enrichment for S1 (orange) or V1 (blue). Point size reflects neighbourhood cell number, edge thickness indicates the number of shared cells between neighbourhoods. (Bottom) Beeswarm plot showing neighbourhood enrichment for S1 or V1. Each point represents a neighbourhood (100–200 cells) with similar expression profiles. The x-axis denotes enrichment (log2 fold change); positive values indicate V1 overrepresentation and negative values S1. Colours indicate significance: grey, not significant; blue, V1-enriched; orange, S1-enriched. CR, Cajal-Retzius cells; Ns, neurons; S1, primary somatosensory cortex; V1, primary visual cortex.

**Figure S2.**
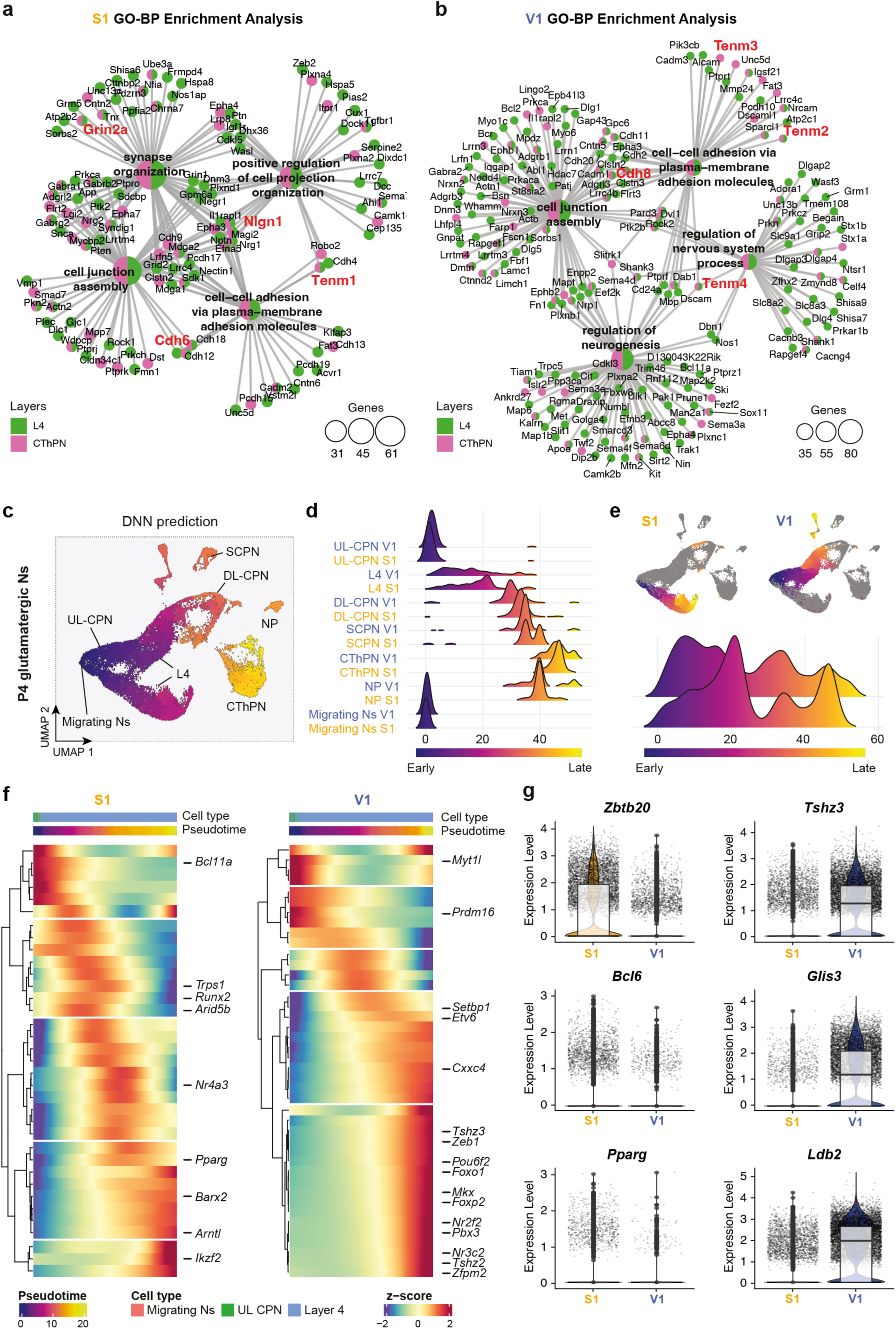
Gene ontology and pseudotime analysis of L4 and CThPN modality-specific genes. **a, b,** Network plots of gene ontology (GO) biological processes (BP) enrichment for modality-specific genes in S1(**a**) and V1(**b**). Node size represents the number of genes associated with each GO term; node colour indicates whether the contributing genes are enriched in L4 (green) or CThPN (pink). **c,** UMAP of inferred pseudotime expression for cortical glutamatergic cell types for S1 and V1. **d,** Density plot showing the cell-type distribution along pseudotime, split by glutamatergic cell-types in V1 and S1**. e,** (Top) Pseudotime trajectories of L4 neurons split by cortical area (S1, left; V1, right). (Bottom) Density plot showing the distribution of sensory modality identity along pseudotime in L4 neurons. **f,** Heatmaps of transcription factors uniquely enriched in S1 (left) and V1 (right), clustered by expression dynamics across pseudotime. **g,** Violin plots showing expression levels of representative transcription factors differentially expressed between S1 and V1. CThPN, corticothalamic projecting neurons; DL CPN, deep-layer cortical projecting neurons; L4, layer 4 neurons; NP, near projecting neurons; Ns, neurons; P, postnatal day; S1, primary somatosensory cortex; SCPN, subcortical projecting neurons; UL CPN, upper-layer cortical projecting neurons; UMAP, uniform manifold approximation and projection; V1, primary visual cortex.

**Figure S3.**
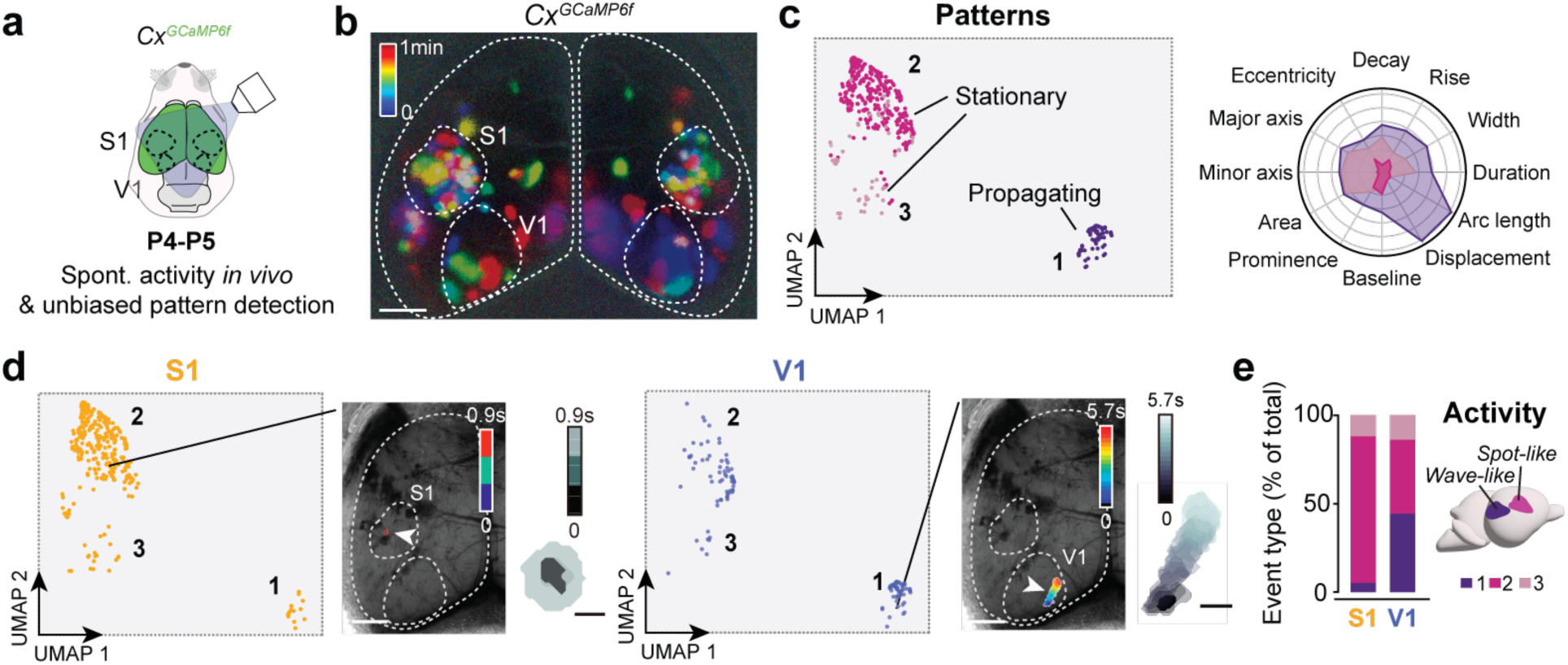
Activity pattern differences between S1 and V1 at P4. **a,** Experimental design for *in vivo* mesoscale calcium imaging. **b,** Temporal colour-coded projection of one minute of spontaneous cortical activity at P4. **c,** (Left) UMAP of all detected events, revealing three distinct clusters: propagating events (cluster 1) and stationary events (clusters 2 and 3). (Right) Radar plot summarizing key properties of each activity profile. **d,** UMAP of calcium events detected separately in S1(left) and V1 (right), with representative examples shown below. **e,** (Left) Quantification of the proportion of each activity profile in S1 and V1 is provided. (Right) Schematic summary of the main findings. S1, primary somatosensory cortex; UMAP, uniform manifold approximation and projection; V1, primary visual cortex. Scale bar, 1000 μm in **b** and **d**, 200 μm in **d** (insets).

**Figure S4.**
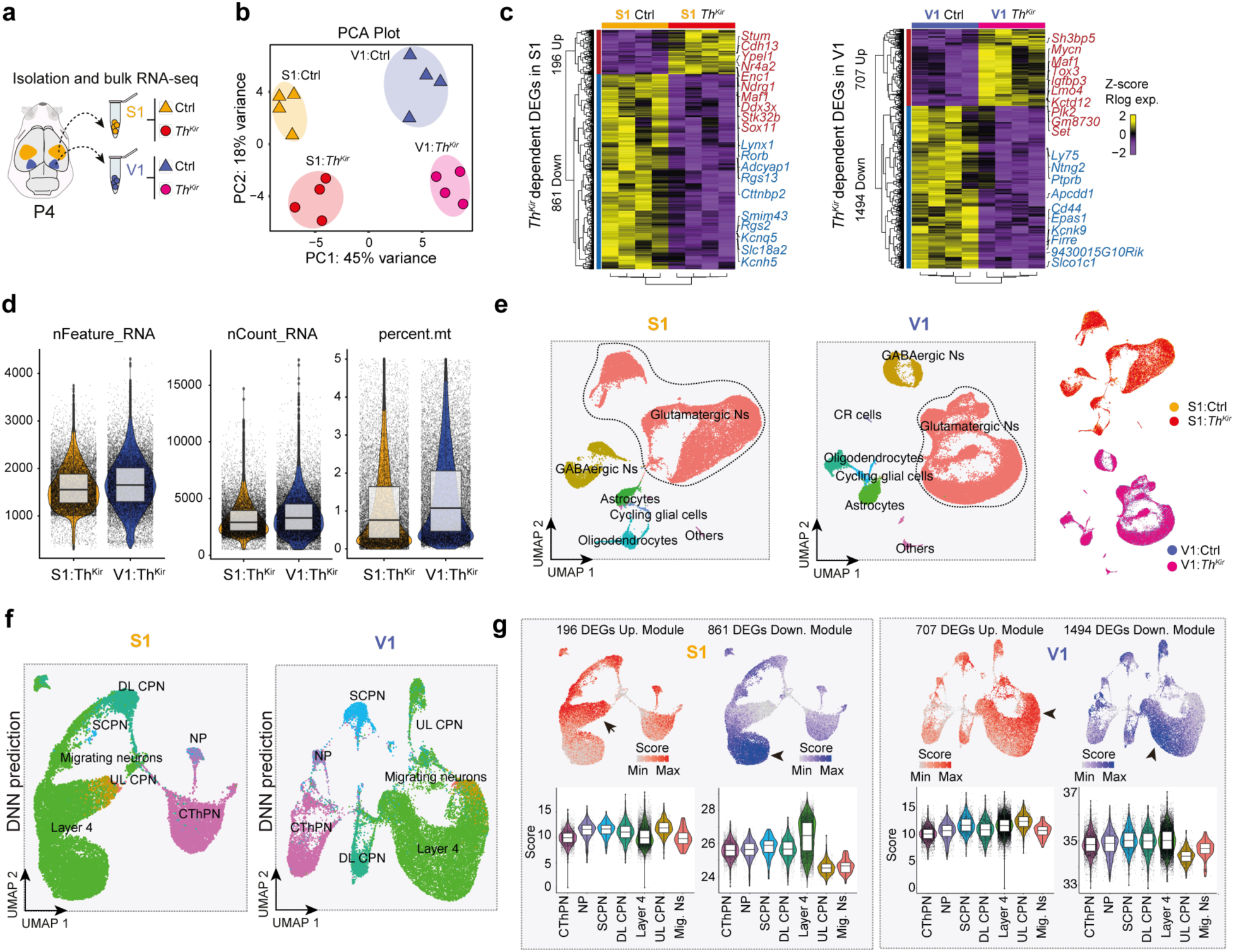
Bulk RNA-seq analysis at P4 showing that *Th^Kir^* primarily affects L4. **a,** Schematic of experimental design. **b,** Principal component analysis (PCA) of control and *Th^Kir^* samples in each cortical region (S1 and V1). **c,** Heatmaps of normalized regularized logarithm (Rlog) z-score expression and unbiased clustering of DEGs between control and *Th^Kir^* samples in S1 (left) and V1 (right). The top 10 DEGs for each condition are highlighted. **d,** Data quality metrics of the *Th^Kir^* snRNA-seq libraries at postnatal day 4 (P4). Violin plots display distributions of total genes detected, total RNA reads, and percentage of mitochondrial gene counts (from left to right) split by cortical regions (S1, orange; V1, blue). **e,** UMAP of the S1 (left) and V1 (middle) snRNA-seq datasets at P4, coloured by identified cell populations. (Right) UMAPs of the same datasets colour-coded by condition (control vs *Th^Kir^*). **f,** UMAPs of glutamatergic neurons from S1 (left) and V1 (right), colour-coded by DNN-predicted cell type. **g,** (Top) UMAPs showing gene module scores for upregulated (red, left) and downregulated (blue, right) DEGs from the S1 bulk RNA-seq dataset, and corresponding plots for V1 (top right). Arrowheads mark L4 neurons. (Bottom) Violin plots of gene module scores across cortical cell types in S1 and V1. CThPN, corticothalamic projecting neurons; DL-CPN, deep-layer cortical projecting neurons; NP, near projecting neurons; Ns, neurons; P, postnatal day; S1, primary somatosensory cortex; SCPN, subcortical projecting neurons; UL-CPN, upper-layer cortical projecting neurons; UMAP, uniform manifold approximation and projection; V1, primary visual cortex.

**Figure S5.**
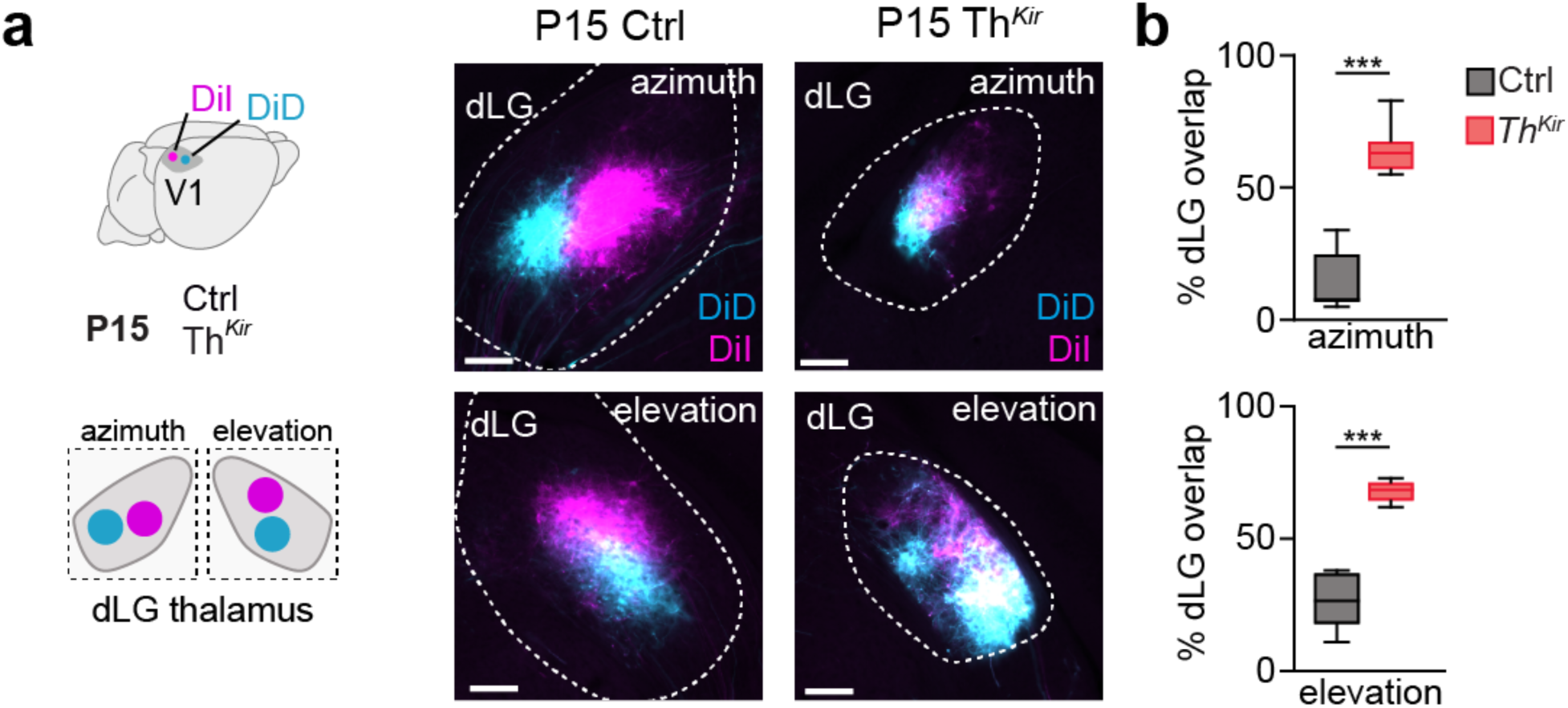
Altered V1 retinotopy in *Th^Kir^*mice. **a,** (Left) Experimental design. (Right) Coronal sections of the dLG showing back-labelling from DiI and DiD injections placed in V1 at P15 in control and *Th^Kir^* mice. **b,** Quantification of azimuth and elevation mapping in control (azimuth, *n* = 6; elevation, *n* = 6) and *Th^Kir^* mice (azimuth, *n* = 8; elevation, *n* = 6). dLG, dorsolateral geniculate nucleus; V1, primary visual cortex. Scale bar 100 μm in **a**.

**Figure S6.**
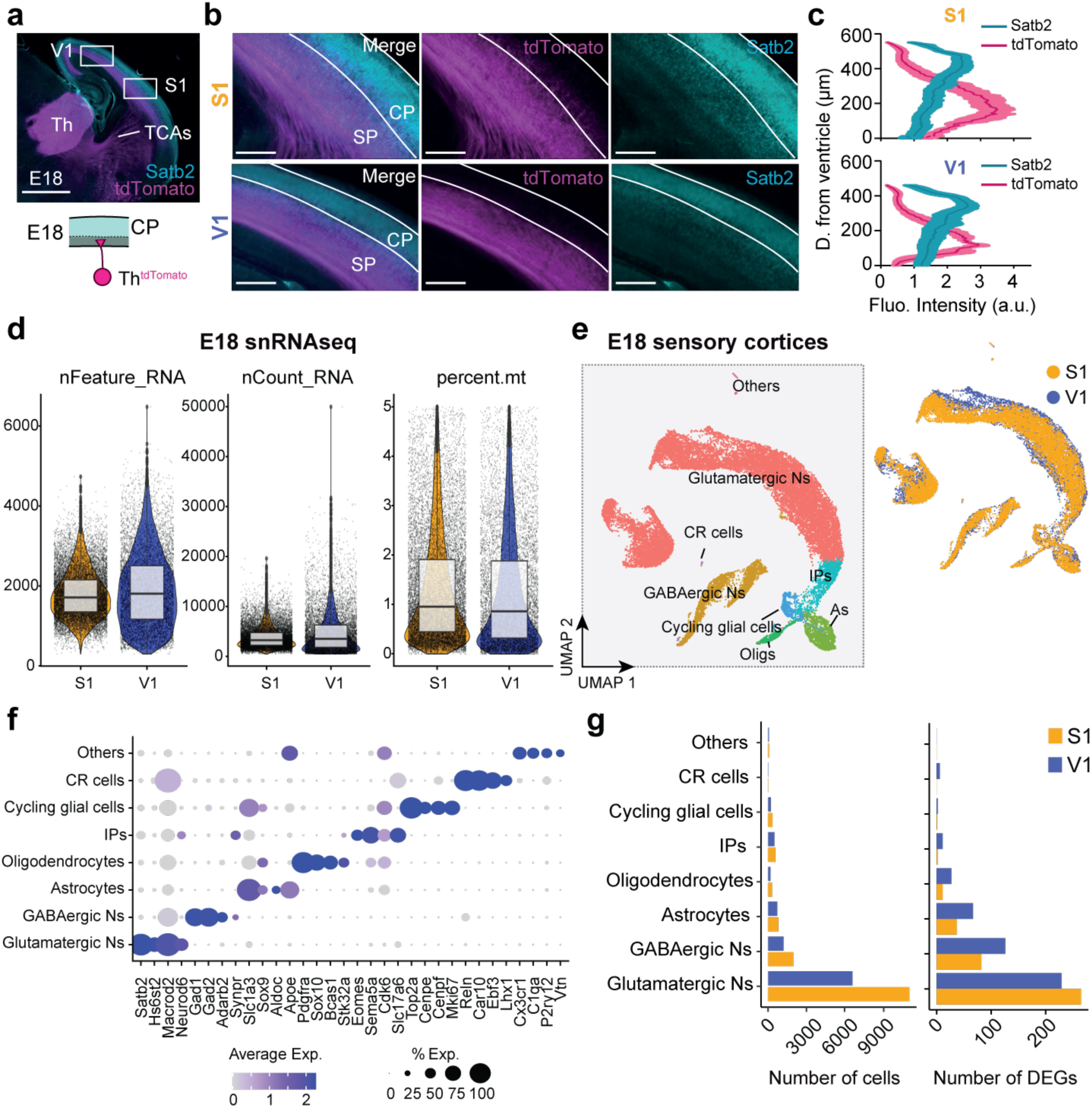
Partial cortical modality specification at embryonic stages. **a,** (Top) Representative 45-degree slice immunostained for Satb2 and tdTomato. (Bottom) Schematic of the experimental design: thalamocortical axon position at E18 was visualized using *Gbx2-cre;tdTomato^fl/fl^* mouse (*Th^tdTomato^*), while the cortical plate was labelled with Satb2 antibody. Gbx2 labels thalamocortical axons, and Satb2 labels upper-layer neurons. **b,** Higher-magnification images of S1 (top) and V1 (bottom) corresponding to the white insets in (**a)**. **c,** Fluorescence intensity profiles of tdTomato and Satb2 across cortical depth in S1 (top) and V1 (bottom) (*n* = 5). **d,** Data quality metrics of the control snRNA-seq libraries at embryonic day 18 (E18). Violin plots display distributions of total genes detected, total RNA reads, and percentage of mitochondrial gene counts (from left to right) split by cortical regions (S1, orange; V1, blue). **e,** (Left) UMAP of the E18 snRNA-seq dataset colour-coded by clusters identified through unsupervised analysis. (Right) UMAP representation colour-coded by the cortical region microdissected. **f,** Dot plot of representative marker genes defining major cell classes at E18. **g,** (Left) Bar plot showing the number of cells assigned to each cortical region (S1, orange; V1, blue). (Right) Bar plot showing the number of differentially expressed genes (DEGs) per region.

**Figure S7.**
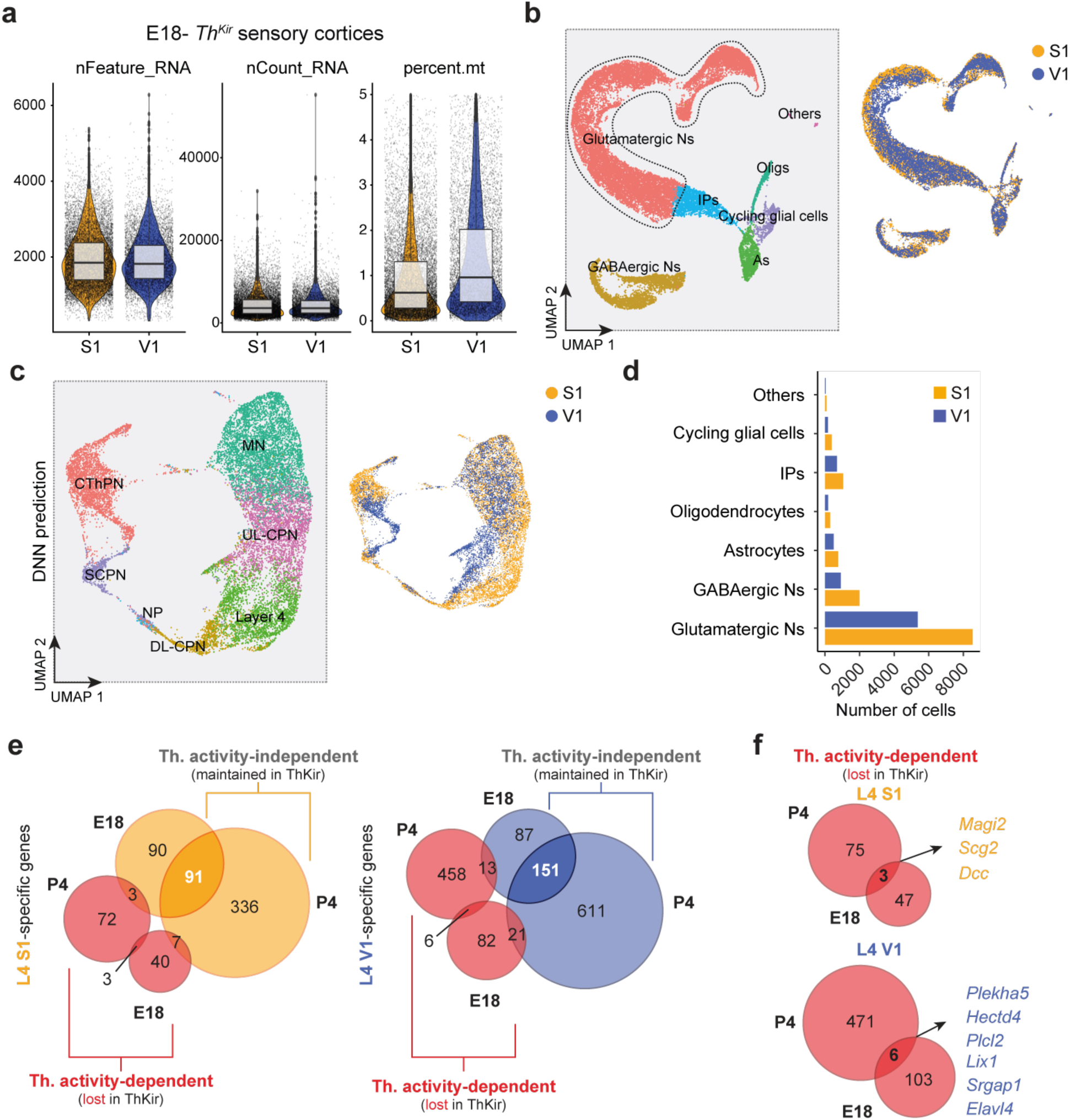
Patterned thalamic inputs selectively regulate identity gene expression in S1 and V1 at E18. **a,** Data quality metrics of the *Th^Kir^* snRNA-seq libraries at embryonic day 18 (E18). Violin plots display distributions of total genes detected, total RNA reads, and percentage of mitochondrial gene counts (from left to right) split by cortical regions (S1, orange; V1, blue). **b,** UMAP of the E18 *Th^Kir^* snRNA-seq dataset coloured by clusters identified through unsupervised analysis (left). UMAP representation coloured by the cortical region microdissected (right). **c,** UMAP visualization of *Th^Kir^* glutamatergic population isolated from the snRNA-seq dataset colour coded by deep neural network (DNN) predicted cell-type (left). UMAP representation coloured by the cortical region microdissected (right). **d,** Bar plot showing the number of cells assigned to each *Th^Kir^* cortical region (S1, orange; V1, blue). **e,** Venn diagrams showing the overlap of differentially expressed genes (DEGs) across developmental stages, cortical areas, and thalamic activity-independent gene sets (S1, left; V1, right). **f**, Venn diagrams showing the overlap of E18 and P4 thalamic activity-dependent DEGs in S1 (top) and V1 (bottom). As, astrocytes; CThPN, corticothalamic projecting neuron; DL-CPN, deep-layer cortical projecting neurons; IPs, intermediate progenitors; L4, layer 4; MN, migrating neurons; NP, near projecting neurons; S1, primary somatosensory cortex; SCPN, subcortical projecting neurons; UL-CPN, upper-layer cortical projecting neurons; UMAP, uniform manifold approximation and projection; V1, primary visual cortex.

**Figure S8.**
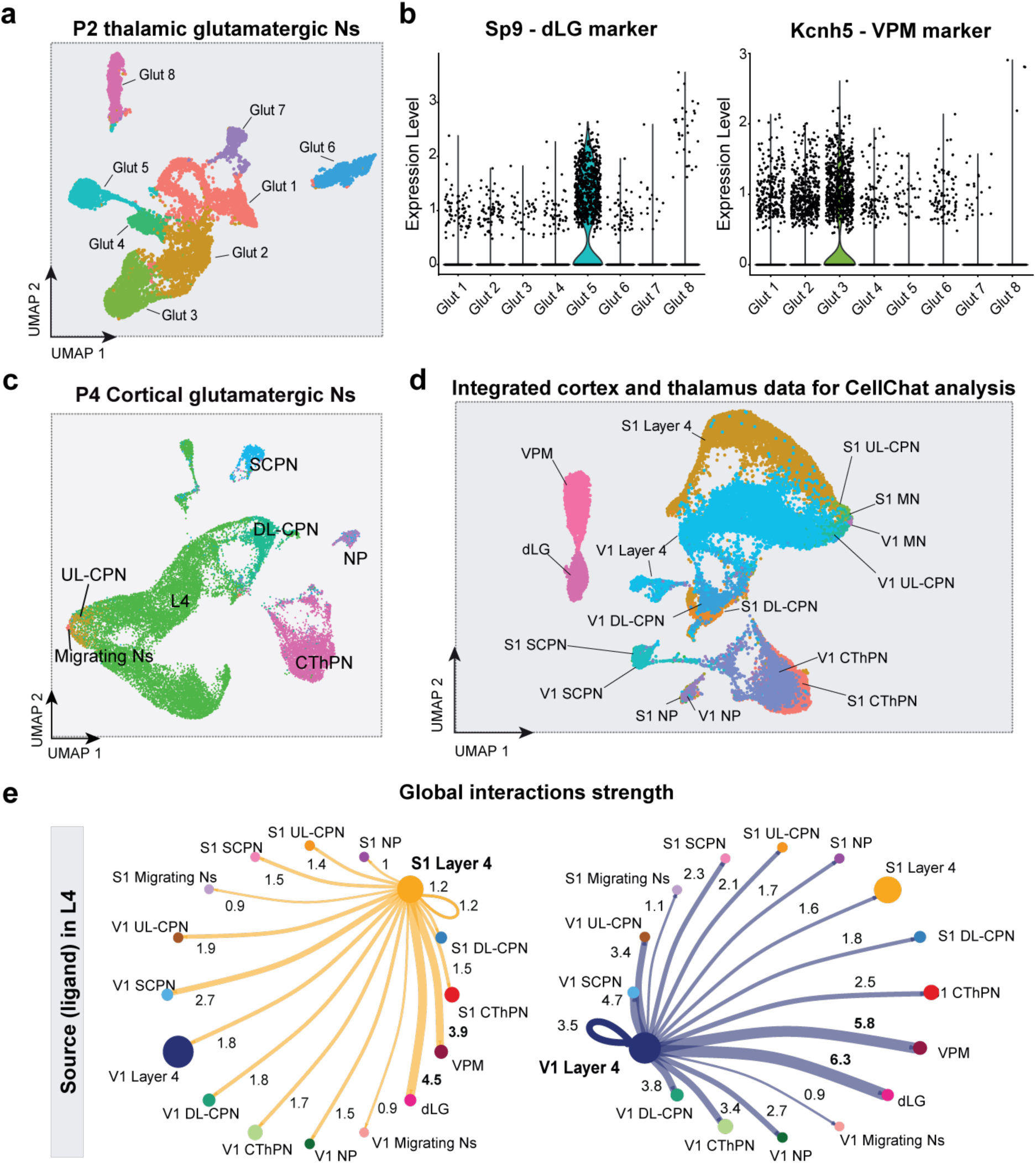
CellChat analysis of intercellular communication between thalamic and cortical neurons. **a,** UMAP of the P2 thalamic snRNA-seq dataset showing identified glutamatergic neuronal clusters. **b,** Violin plots displaying representative marker genes for dLG (left) and VPM (right) neurons. **c,** UMAP of the P4 cortical snRNA-seq dataset annotated with major cortical neuronal types. **d,** UMAP of the combined dataset integrating P4 cortical neurons with P2 thalamic dLG and VPM neurons used for CellChat analysis. **e,** CellChat-inferred outgoing interactions from L4 neurons in S1 (left) and V1 (right). Circle size indicates the number of cells per group; edge width and labels represent communication probability. CThPN, corticothalamic projecting neurons; DL-CPN, deep-layer cortical projecting neurons; dLG, dorsolateral geniculate nucleus; L4, layer 4 neurons; NP, near-projecting neurons; Ns, neurons; P, postnatal day; S1, primary somatosensory cortex; SCPN, subcortical projecting neurons; UL-CPN, upper-layer cortical projecting neurons; UMAP, uniform manifold approximation and projection; V1, primary visual cortex; VPM, ventral posteromedial nucleus.

**Figure S9.**
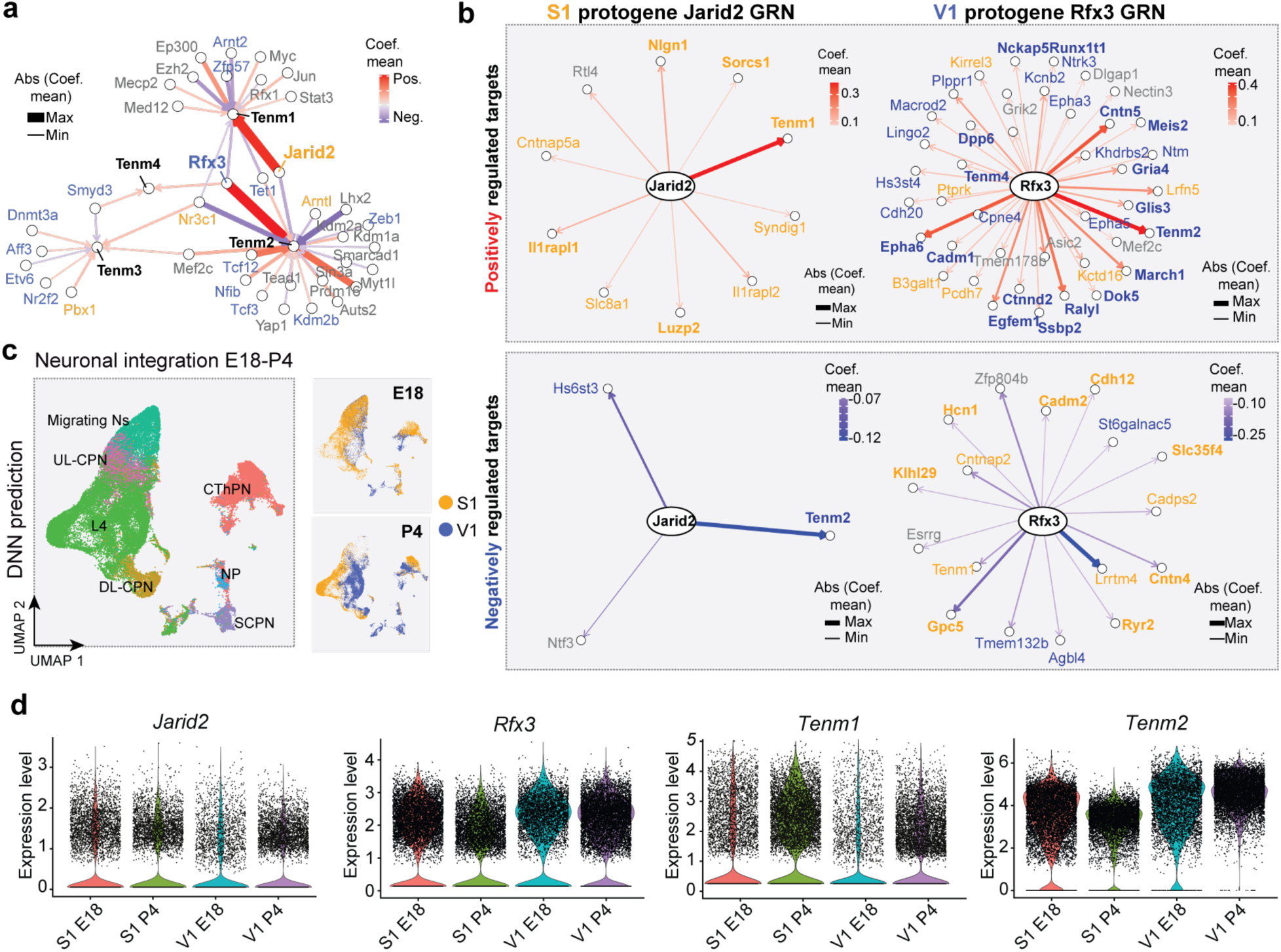
Gene regulatory network analysis identifies candidate transcriptional regulators of modality-specific programs. **a,** Node–edge diagram of predicted upstream transcriptional regulators of teneurins. *Jarid2* emerges as the strongest predicted regulator of *Tenm1* (S1-enriched), while *Rfx3* is the strongest predicted regulator of *Tenm2* (V1-enriched). **b,** Gene regulatory networks (GRNs) for *Jarid2* and *Rfx3*. Target genes are coloured according to area specificity (orange, S1-upregulated; blue, V1-upregulated) at P4. Protogenes are shown in bold. **c,** UMAP visualization of integrated glutamatergic neurons across E18–P4 (left). Corresponding UMAPs colour-coded by cortical region (S1, orange; V1, blue) at E18 (top right) and P4 (bottom right). **d,** Violin plots showing expression of selected genes across developmental stages and modalities. CThPN, corticothalamic projecting neurons; DL-CPN, deep-layer cortical projecting neurons; L4, layer 4 neurons; NP, near projecting neurons; Ns, neurons; P, postnatal day; S1, primary somatosensory cortex; SCPN, subcortical projecting neurons; UL-CPN, upper-layer cortical projecting neurons; UMAP, uniform manifold approximation and projection; V1, primary visual cortex.

**Figure S10.**
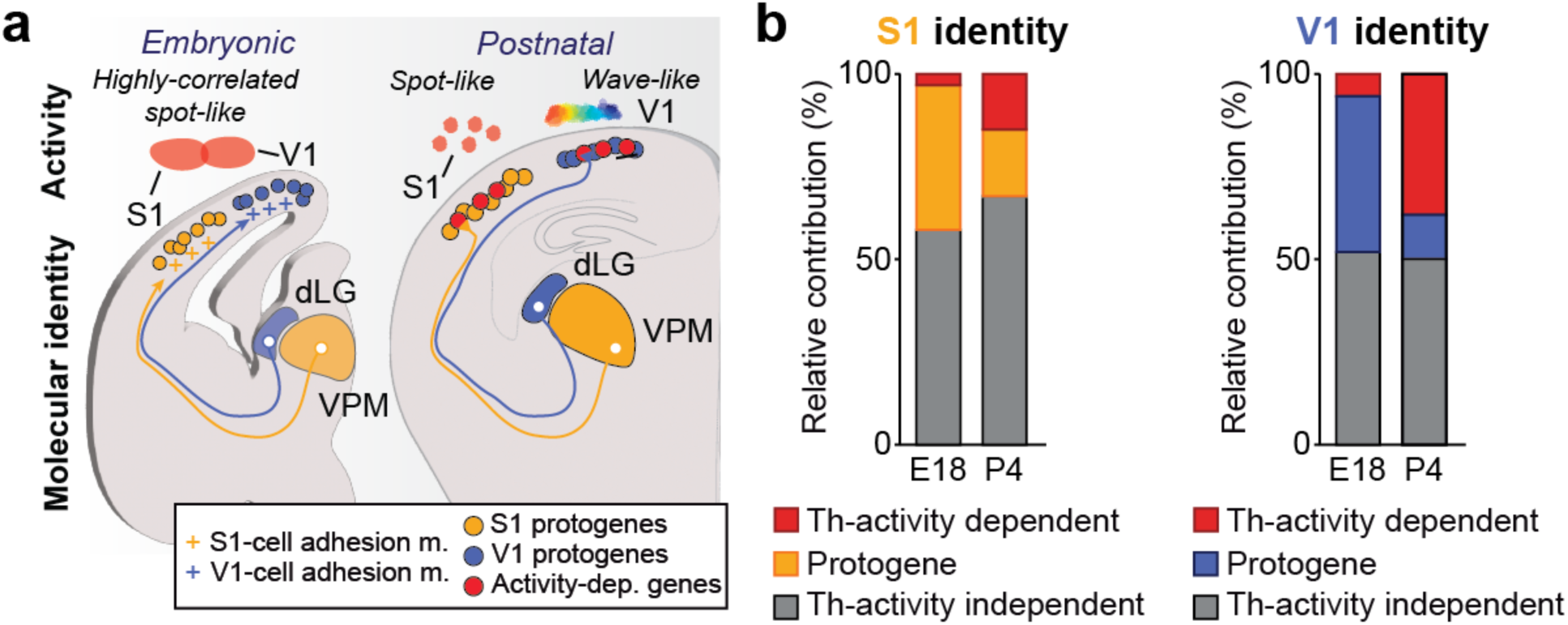
Schematic summary of molecular and activity-dependent contributors to cortical modality specification. **a,** Conceptual model summarizing developmental contributions to cortical sensory identity. At embryonic stages, cortex-intrinsic expression of cell-adhesion molecules forms early, region-specific molecular signatures that may help align thalamocortical projections through homophilic or heterophilic interactions. Incoming thalamic input from modality-specific nuclei contributes additional transcriptional refinement, supporting the emergence of distinct sensory identities. By postnatal stages, these combined intrinsic and activity-dependent influences give rise to functionally divergent cortical modalities, each displaying characteristic patterns of spontaneous activity. **b,** Proportion of S1- and V1-identity genes classified as thalamic activity-dependent, activity-independent, or protogenes at E18 and P4. Thalamic activity shows a minor contribution to modality-specific transcriptional identity at embryonic stages but a stronger contribution at postnatal stages in both sensory cortices. dLG, dorsolateral geniculate nucleus; VPM, ventral posteromedial nucleus; S1, primary somatosensory cortex; V1, primary visual cortex.

